# GolpHCat (TMEM87A): a unique voltage-gated and pH-sensitive cation channel in the Golgi

**DOI:** 10.1101/2023.01.03.522543

**Authors:** Hyunji Kang, Heejin Jeong, Ah-reum Han, Wuhyun Koh, Jung Moo Lee, Heeyoung Jo, Hayeon Lee, Mridula Bhalla, Woo Suk Roh, Hyun Jun Jang, Boyoung Lee, Ho Min Kim, Hyun Joo An, C. Justin Lee

## Abstract

The Golgi apparatus is a critical intracellular organelle that is responsible for modifying, packaging, and transporting proteins to their destinations. Golgi homeostasis involving the acidic pH, ion concentration, and membrane potential, is critical for proper functions and morphology of the Golgi. Although transporters and anion channels that contribute to Golgi homeostasis have been identified, the molecular identity of cation channels remains unknown. Here we identify TMEM87A as a novel Golgi-resident cation channel that contributes to pH homeostasis and rename it as GolpHCat (**Gol**gi **pH**-sensitive **Cat**ion channel). The genetic ablation of GolpHCat exhibits an impaired resting pH in the Golgi. Heterologously expressed GolpHCat displays voltage- and pH-dependent, non-selective cationic, and inwardly rectifying currents, with potent inhibition by gluconate. Furthermore, reconstitution of purified GolpHCat in liposomes generates functional channel activities with unique voltage-dependent gating and ion permeation. GolpHCat is expressed in various cell types such as neurons and astrocytes in the brain. In the hippocampus, GolpHCat-knockout mice show dilated Golgi morphology and altered glycosylation and protein trafficking, leading to impaired spatial memory with significantly reduced long-term potentiation. We elucidate that GolpHCat, by maintaining Golgi membrane potential, regulates ionic and osmotic homeostasis, protein glycosylation/trafficking, and brain functions. Our results propose a new molecular target for Golgi-related diseases and cognitive impairment.

## Main

Golgi homeostasis is critical for proper Golgi morphology and functions such as modifying, packaging, and transporting proteins. Impaired Golgi homeostasis causes altered protein trafficking and glycosylation^1,2^, and induces morphological changes in the Golgi, such as fragmentation, which is associated with diseases involving neurodegenerative diseases^3–5^. Golgi homeostasis is proposed to be regulated by an ATP-mediated proton pump, proton leak exchanger, proton leak channel, anion channel, and cation channel^6^. The ATP-mediated proton pump and protein leak exchanger have been identified as V-ATPase^7^ and NHE7/8^8,9^, respectively. The anion channel regulating Golgi pH and morphology has so far been identified as GPHR^10^, which maintains normal neuronal morphology and circuitry^11^. However, the molecular identity and functions of Golgi-resident cation channels in the brain remain elusive.

Potential candidate for Golgi-resident cation channel could be one of the TMEMs which are transmembrane proteins with unknown functions expressed in the plasma membrane or intracellular organelle membrane. Over the past decade, several TMEMs and their cryogenic electron microscopy (cryo-EM) structures have been extensively investigated and found to be functional ion channels^12–21^. Recently, a TMEM with generic name of TMEM87A has been reported as a mainly Golgi-localized protein involved in protein trafficking^22,23^. However, it has not been determined whether TMEM87A is an ion channel or not. Indeed, TMEM87A has been recently proposed as either a pore-forming or auxiliary subunit for a mechanosensitive ion channel^24^, which has been immediately contradicted by the lack of mechanosensitive channel activities in TMEM87A reconstituted proteoliposome^25^.

### TMEM87A is localized in the Golgi apparatus and contributes to Golgi pH homeostasis

To investigate whether TMEM87A is an ion channel or not, we first analyzed the protein sequence of TMEM87A and found that TMEM87A contains a GYG sequence which is a signature selectivity filter of classical K^+^ channels^26^ (Fig. 1a), raising a possibility that TMEM87A may be a cation channel. *TMEM 87A* gene encodes a 63-kDa protein with a predicted N-terminal Golgi signal sequence and seven transmembrane (TM) domains (Fig. 1b, Extended Data Fig. 1a-b). In humans, *TMEM87A* gene encodes for three isoforms: isoform 1 is full-length with a predicted Golgi signal sequence and TMs, isoform 2 has no TMs, and isoform 3 has no predicted Golgi signal sequence (Fig. 1c, Extended Data Fig. 1a). According to the brain RNA-seq database, *TMEM87A* is highly expressed in the neurons and astrocytes of the brain^27,28^. Thus, based on bioinformatics, TMEM87A is a potential candidate for Golgi-resident cation channel in the brain.

**Figure 1.**
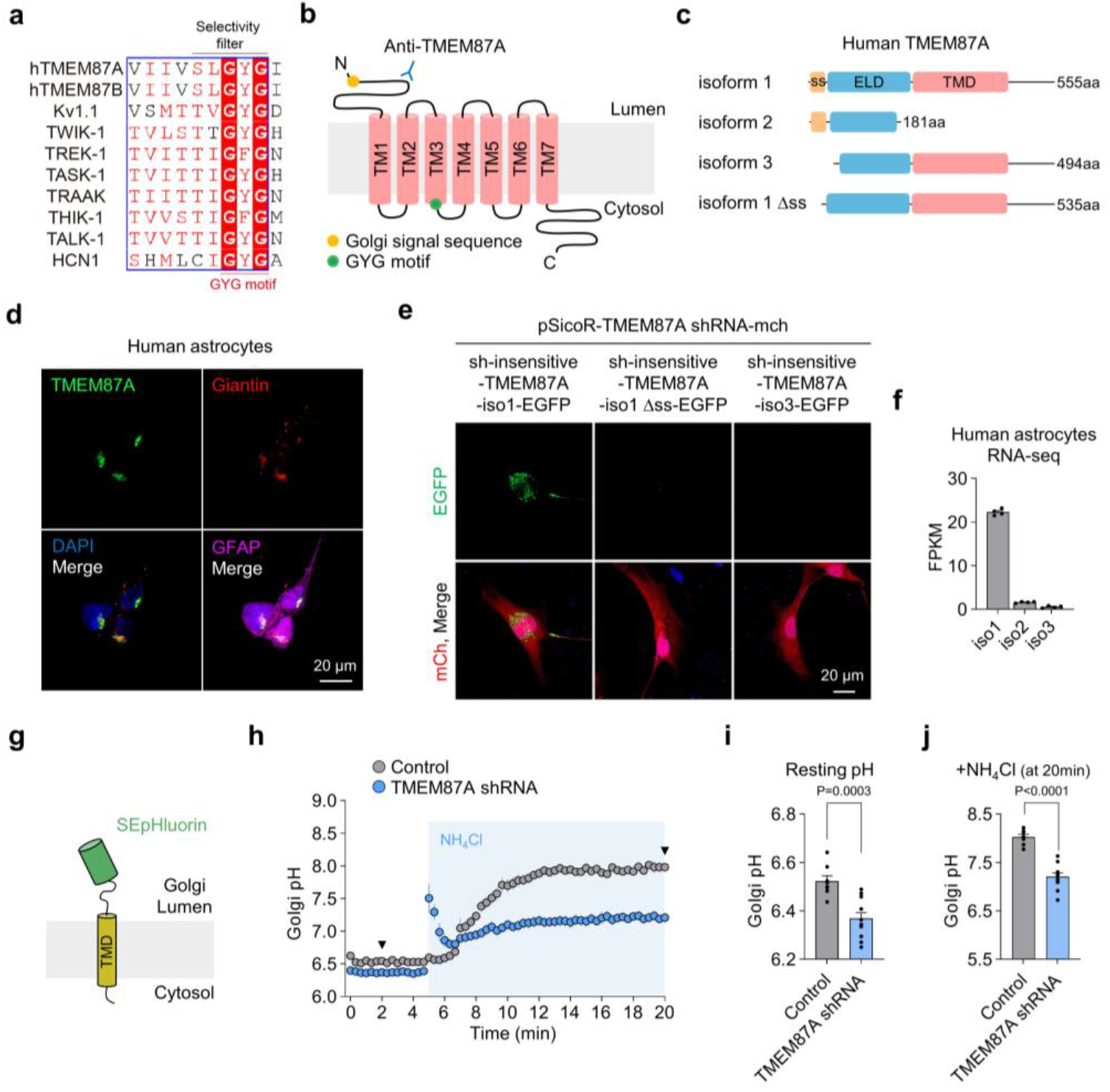
TMEM87A is localized in the Golgi apparatus and contributes to Golgi pH homeostasis. **a,** Partial sequence alignment for the selectivity filter of K^+^ channel family with hTMEM87A/B. **b,** A predicted seven-transmembrane (TM) topology of TMEM87A within the Golgi lumen. Anti-TMEM87A indicates epitope for TMEM87A antibody. Green circle indicates a GYG motif sequence. **c,** Proteins of human TMEM87A. ELD, extracellular domain; TMD, transmembrane domain; ss, signal sequence; isoform 1 Δss, ss deleted isoform 1. **d,** Colocalization of TMEM87A with the Giantin, Golgi marker, in cultured human astrocytes. **e,** Localization of C-terminal EGFP-tagged shRNA-insensitive TMEM87A isoforms under the gene-silencing of endogenous TMEM87A in cultured human astrocytes. **f,** Transcriptome analysis of TMEM87A isoforms using RNA-seq in cultured human astrocytes. **g,** Schematic diagram of used construct ‘B4GALT1-SEpHluorin’ for measuring pH of Golgi lumen. **h-j,** Comparison of resting Golgi luminal pH and buffer capacities under the absence and presence of 100 mM NH4Cl for 15 min in Control (n=8), and TMEM87A shRNA transfected (n=11) cultured human astrocytes expressing B4GALT1-SEpHluorin. Arrows indicate time points in (**i,j**). **h,** Golgi pH before and after treating the 100 mM NH_4_Cl. **i,** Golgi pH values before treating the NH_4_Cl from (**h**). **j,** Golgi pH values after treating the 100 mM NH4Cl for 15 min from (**h**). Data are presented as the mean ± SEM. Student two-tailed unpaired t-test in (**i, j)**.

To examine the protein expression of TMEM87A in Golgi, we performed immunocytochemistry (ICC) with antibodies against TMEM87A and Giantin as a Golgi marker in cultured human astrocytes (Fig. 1d). We found that TMEM87A was highly colocalized with Giantin (Pearson’s coefficient R: 0.59 ± 0.04) in cultured human astrocytes (Fig. 1d), indicating that TMEM87A is mainly localized in Golgi. To investigate whether the predicted Golgi signal sequence is indeed responsible for Golgi localization, we overexpressed full DNA sequence of each isoform 1 and predicted Golgi signal sequence deleted isoform1 (isoform 1 Δss), and predicted Golgi signal sequence lacking isoform 3 (Fig.1c) in cultured human astrocytes. To minimize the contribution of endogenous wild-type TMEM87A, we developed TMEM87A shRNA and co-transfected with each shRNA-insensitive EGFP-tagged TMEM87A isoform in cultured human astrocytes (Fig 1e). We observed that isoform 1 was localized in Golgi, but isoform 1 Δss and isoform 3 were not observed in Golgi (Fig. 1e), indicating that the predicted N-terminal signal sequence is required for Golgi localization of TMEM87A. Furthermore, we employed the Next Generation RNA-sequencing (RNA-seq) to examine the expression level of isoforms in cultured human astrocytes (Fig. 1f). We found that TMEM87A isoform 1 showed the greatest proportion of total TMEM87A expression (Fig. 1f). Taken together, these results indicate that the major form of TMEM87A, isoform 1 is localized in Golgi due to N-terminal signal sequence in cultured human astrocytes.

To examine whether TMEM87A contributes to Golgi pH, we developed and expressed a Golgi luminal-targeting pH sensor construct, B4GALT1-SEpHluorin^29^, for real-time imaging of pH in cultured human astrocytes (Fig. 1g, Extended Data Fig. 2a). We found that gene-silencing of TMEM87A by TMEM87A shRNA led to more acidic resting Golgi pH compared to the control (Fig. 1h, i, Extended Data Fig.2b, c). Furthermore, the Golgi pH buffer capacity, as measured by a change in pH upon 100 mM NH4Cl application, was significantly lower in TMEM87A shRNA transfected cells (Fig. 1h, j, Extended Data Fig.2b, c), indicating that TMEM87A contributes to Golgi pH buffer capacity. Taken together, these results indicate that TMEM87A, a candidate cation channel, localizes in Golgi and contributes to Golgi pH homeostasis.

### TMEM87A is a voltage- and pH-dependent, non-selective, inwardly rectifying cation channel

Next, to investigate whether TMEM87A mediates current in the heterologous expression system, we transfected human TMEM87A in the CHO-K1 cells and recorded whole-cell currents under voltage-clamp (Fig. 2a). Although the native TMEM87A is mainly localized in Golgi of human astrocytes, we observed that the EGFP-tagged TMEM87A under heterologous overexpression system was found not only in Golgi but also in plasma membrane (Fig. 2a). Firstly, we measured voltage-dependent membrane current under the voltage-ramp protocol ranging from −150 mV to +100 mV with 140 mM NaCl containing external solution and 130 mM K-gluconate containing internal solution (Fig. 2b). TMEM87A WT-mediated current displayed a non-linear current-voltage (I-V) relationship with a reversal potential near −7.7 mV and pronounced inward-rectification near −150 mV (Fig. 2b-d). The average rectification index value of TMEM87A WT from +100 mV to −150 mV was 2.7 ± 0.3 (Fig. 2e). In contrast to TMEM87A WT, both outward and inward currents were completely abolished in TMEM87A-AAA (GYG318~320AAA) mutant transfected cells (Fig. 2b-e). We performed surface biotinylation assay and confirmed the surface expression of both TMEM87A WT and AAA mutant forms (Extended Data Fig. 3a). These results indicate that GYG sequence in TMEM87A might have a critical role in mediating current. Furthermore, we recorded currents with the voltage-step pulses from +100 mV to −150 mV and found that TMEM87A-mediated current displayed a voltage-dependent inward rectification with no time- and voltage-dependent inactivation (Fig. 2f). Taken together, TMEM87A mediates voltage-dependent inwardly rectifying membrane currents in the heterologous expression system.

**Figure 2.**
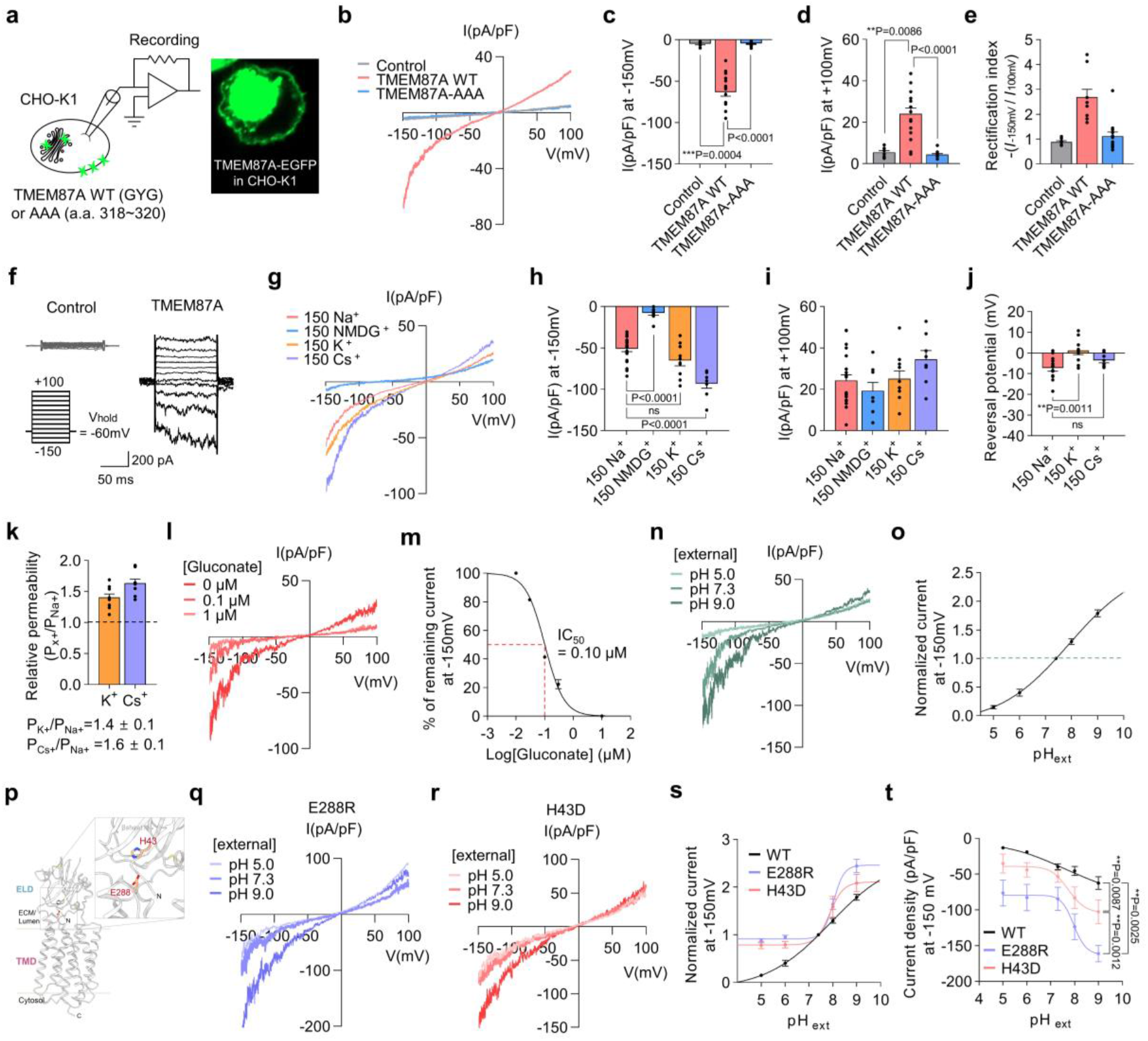
TMEM87A mediates voltage- and pH-dependent, inwardly rectifying cationic currents. **a,** Schematic diagram of whole-cell patch-clamp recording from TMEM87A WT- or TMEM87A-AAA (a.a. 318~320)-IRES2-DsRed transfected CHO-K1 cell. Inset, fluorescence image of EGFP-tagged TMEM87A. **b,** Averaged I-V relationship from Control (gray; n=7), TMEM87A WT (pink; n=16), or TMEM87A-AAA transfected (blue; n=12) cells under voltage ramp protocol (from +100 mV to −150 mV). **c-d,** Current densities measured at −150 mV (**c**) and at +100 mV (**d**). **e,** Rectification index calculated as the absolute ratio of amplitude at −150 mV over at +100 mV. **f,** Representative currents from Control and TMEM87A WT transfected cells under voltage step protocol (from +100 mV to −150 mV, 25 mV step). **g,** Averaged I-V relationship from TMEM87A WT transfected cells with bath solutions containing Na^+^ (n=21), NMDG^+^ (n=8), K^+^ (n=10), or Cs^+^ (n=8). **h-i,** Current densities measured at −150 mV (**h**) and +100 mV (**i**). **j,** Reversal potentials in Na^+^, K^+^, or Cs^+^-containing bath solutions. **k,** Relative permeability ratio of Na^+^ to K^+^ (P_K+_/P_Na+_) or Cs^+^ (P_Cs_+/P_N+_). **l,** Representative I-V relationship from TMEM87A WT transfected cell with or without gluconate in bath solution. **m,** Dose-response curve for percentage currents at −150 mV for gluconate (0, 0.01, 0.03, 0.1, 0.3, and 10 μM) (n=5). **n,** Representative I-V relationship from TMEM87A WT transfected cell under various pH. **o,** Normalized currents at −150 mV under various pH, normalized to current at pH 7.3 (pH 5, 6, 7.3, 8, and 9) (n=7). **p,** Predicted key interacting residues which are shown as orange sticks and labeled in the cryo-EM structure of hTMEM87A (PDB ID: 8HSI). **q-r,** Representative I-V relationship from hTMEM87A-E288R mutant (**q**) or H43D mutant (**r**) transfected cells under various pH. **s,** Normalized currents at −150 mV of hTMEM87A WT (n=7), E288R (n=5), or H43D (n=4). **t,** Current densities measured at −150mV under various pH. Data are presented as the mean ± SEM. One-way ANOVA followed by Dunn’s multiple comparisons test in (**c,d)** or Dunnett’s multiple comparisons test in (**h,i,j,t)**.

To investigate whether TMEM87A-mediated inward current is carried by Na^+^ ion, we replaced Na^+^ with N-methyl-D-glucamine (NMDG) (Fig. 2g). The inward current was mostly abolished, suggesting that TMEM87A might be a Na^+^-permeable cation channel (Fig. 2g-i). To determine the relative permeability ratio of TMEM87A-mediated currents to different cations such as K^+^ and Cs^+^, we replaced Na^+^ with K^+^ or Cs^+^ (Fig. 2g-i) and found that reversal potentials were slightly shifted to more positive potentials; from Na^+^ (−7.7 ± 1.5 mV) to K^+^ (0.5 ± 1.6 mV) and Cs^+^ (−3.5 ± 1.1 mV) (Fig. 2j). Using a modified Goldman-Hodgkin-Katz equation, we calculated the permeability ratios to be P_K+_/P_Na+_ = 1.4 ± 0.1 and P_Cs+_/P_Na+_ = 1.6 ± 0.1 (Fig. 2k), suggesting that TMEM87A might be a non-selective cation channel with a slightly higher permeability to K^+^ and Cs^+^ compared to Na^+^. We further confirmed that TMEM87A-mediated current was not carried by Cl^-^ (Extended Data Fig. 3b, c). Taken together, these results provide a series of evidence that TMEM87A might be a non-selective cation channel.

To investigate the pharmacological properties of TMEM87A-mediated currents, we tested the inhibitory effect of gadolinium (Gd^3+^), which is well known non-selective cation channel blocker. We found that Gd^3+^ effectively blocked TMEM87A-mediated currents in dose-dependent manner with a half maximal inhibition (IC_50_) of 0.91 μM (Extended Data Fig. 3d, e). Like human TMEM87A, mouse TMEM87A mediated similar magnitude of membrane currents with similar gadolinium sensitivity (Extended Data Fig. 3f, g). While we were substituting Cl^-^ with various large anions, we serendipitously discovered that gluconate potently blocked TMEM87A-mediated currents with IC_50_ of 0.10 μM (Fig. 2l, m).

To investigate if TMEM87A-mediated current is sensitive to pH, we recorded currents at different pH bath solutions (Fig. 2n). We found that TMEM87A-mediated inward current was significantly reduced in acidic pH, whereas it was significantly enhanced in basic pH (Fig. 2o), indicating that TMEM87A-mediated current is pH-sensitive. Despite its pH-sensitivity, TMEM87A showed negligible permeability to H^+^ (Extended Data Fig. 3h, i). It has been reported that a hydrogen bond or salt bridge between histidine and glutamate residues leads to pH-dependent conformational changes in oligomycin sensitivity conferral protein (OSCP) subunit of mitochondrial F1FO (F)-ATP synthase^30^. Based on the structure predicted by AlphaFold2^31,32^ (Fig. 2p), we investigated whether the proximately located E288 and H43 are the key residues for pH-sensing mechanism of TMEM87A-mediated currents (Fig. 2q-r), as both residues are located in the extracellular domain of the protein (corresponding to Golgi lumen) and H43 becomes protonated at acidic pH. We then recorded whole-cell currents at different pH bath solutions mediated by E288R or H43D mutations thereby reversing the charge on the respective amino acids (Fig. 2q-r). We found that TMEM87A-E288R and H43D-mediated currents at −150 mV were significantly larger than that of WT, and the pH-sensitivity was eliminated especially in acidic pH’s (Fig. 2s-t). These results suggest that TMEM87A-mediated currents may be sensitive to pH due to the electrostatic interaction between E288 and H43. Taken together, these results support that TMEM87A might be a novel voltage- and pH-dependent, non-selective, inwardly rectifying cation channel.

### GolpHCat (TMEM87A) is a *bona fide* ion channel

To investigate whether TMEM87A is a *bona fide* functional ion channel, we purified and reconstituted EGFP-tagged TMEM87A proteins in liposomes (proteoliposome), and then performed blister-attached patch recording for single channel activities with 200 mM KCl symmetric solutions (Fig 3a). TMEM87A in proteoliposome displayed conspicuous stochastic single channel openings at positive or negative holding potentials from +90 mV to −150 mV, but not at 0 mV (Fig. 3b). Detailed analysis showed that the channel’s open probability (Po) starts from Po = 0 at 0 mV and increases non-linearly at both negative and positive potentials, with maximum Po ≈ 0.6 at +90 mV and Po ≈ 0.3 at −150 mV (Fig. 3c). These results indicate that the purified TMEM87A is a voltage-dependent channel with activation voltages at both negative and positive potentials and with much higher Po at positive potentials (6-fold higher Po at +90 mV compared to Po at −90 mV; Fig. 3c). The analysis of amplitude showed a weak inward rectification (Fig. 3d), which is in marked contrast to the strong inward rectification in the whole-cell patch results (Fig. 2b). However, when we multiplied the unitary current and open probability at each holding potential, we then obtained a strongly rectifying I-V relationship (Fig. 3e), implying that the whole-cell patch results were genuinely originated from the ensemble average of single channel activities of TMEM87A. Interestingly, TMEM87A showed no subconductance opening at negative potentials including −150 mV, whereas there were numerous subconductance-level openings (O_sub_) at +90 mV (Fig. 3b, f). In addition, we found that the single channel conductance was gradually increased at negative potentials, whereas it was gradually decreased at positive potentials (Fig. 3b). These results indicate that the voltage-dependent gating and ion permeation of TMEM87A are profoundly different at positive and negative potentials. The open- and closed-time distribution plots showed that TMEM87A frequently opened and closed for brief periods at +90 mV with time constants, τ_open_ = 26 ms and τ_close_ = 41 ms (Fig. g, h), whereas it less frequently opened and closed for longer periods at −150 mV with time constants, Topen = 421 ms and τ_close_ = 505 ms (Fig. 3i, j), again indicating a profoundly different channel gating at positive and negative potentials. Using the Liquid Chromatography mass spectrometry (LC-MS), in the purified TMEM87A solutions, we found negligible amount of other pore-forming ion channel proteins (Extended Data Fig. 4a, b, Table 1) which might confound our conclusion that TMEM87A is a *bona fide* ion channel. Taken together, these results provide direct evidence that TMEM87A is a voltage-gated cation channel. Based on our results, we renamed TMEM87A as GolpHCat to represent a novel **Gol**gi-resident, **pH**-sensitive, **Cat**ion channel.

**Figure 3.**
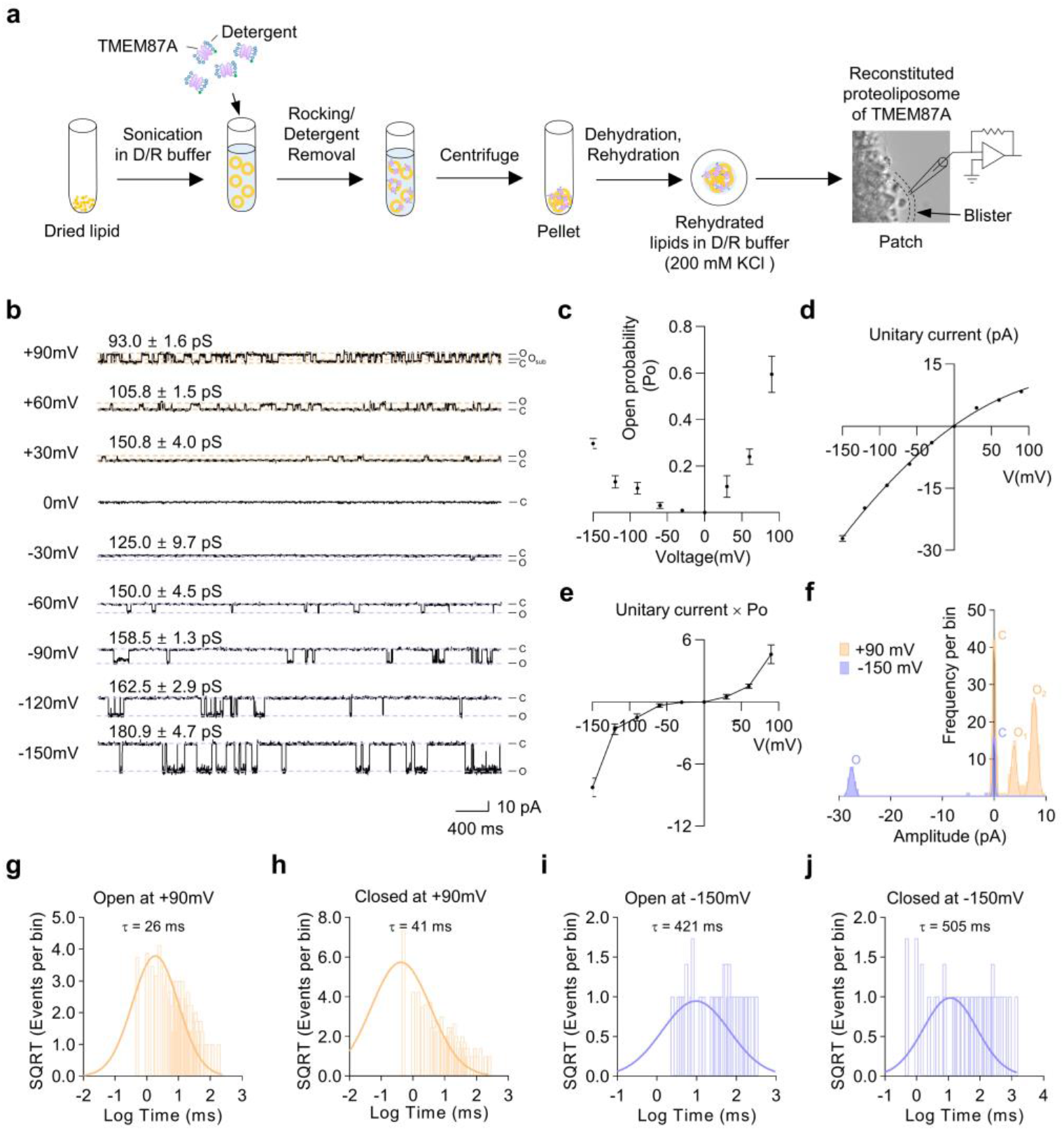
TMEM87A is *bona fide* functional ion channel in proteoliposome. **a,** Schematic diagram showing the procedure for reconstitution of TMEM87A into liposome with dehydration/rehydration method for single-channel recording. **b,** Representative spontaneous single channel currents from the reconstituted proteoliposome of TMEM87A under voltage steps (from +90 mV to −150 mV, 30 mV step) from the same patch condition. The single channel conductance values were calculated at each holding potential. **c,** Voltage dependent channel opening probability (Po) of TMEM87A at each holding potential. **d,** I-V relationship of TMEM87A single channel unitary current activities. Data was fitted with a polynomial. **e,** The unitary current × open probability-voltage relationship of TMEM87A from (**c,d**). **f,** Amplitude histogram of TMEM87A single channel unitary current activities with open (O) and closed (C) states at +90 mV (orange) and −150 mV (purple) from (**b**). The distribution data are fitted with a sum of two Gaussians at each holding potential. **g-h,** Dwell time histograms of TMEM87A single channel unitary current activities at +90 mV for open (**g**) and closed (**h**) states (n=3). The data are fitted with log-normal distribution. **i-j,** Dwell time histograms of TMEM87A single channel activity at −150 mV for open (**i**) and closed (**j**) states (n=3). The data are fitted with log-normal distribution.

**Table 1.**
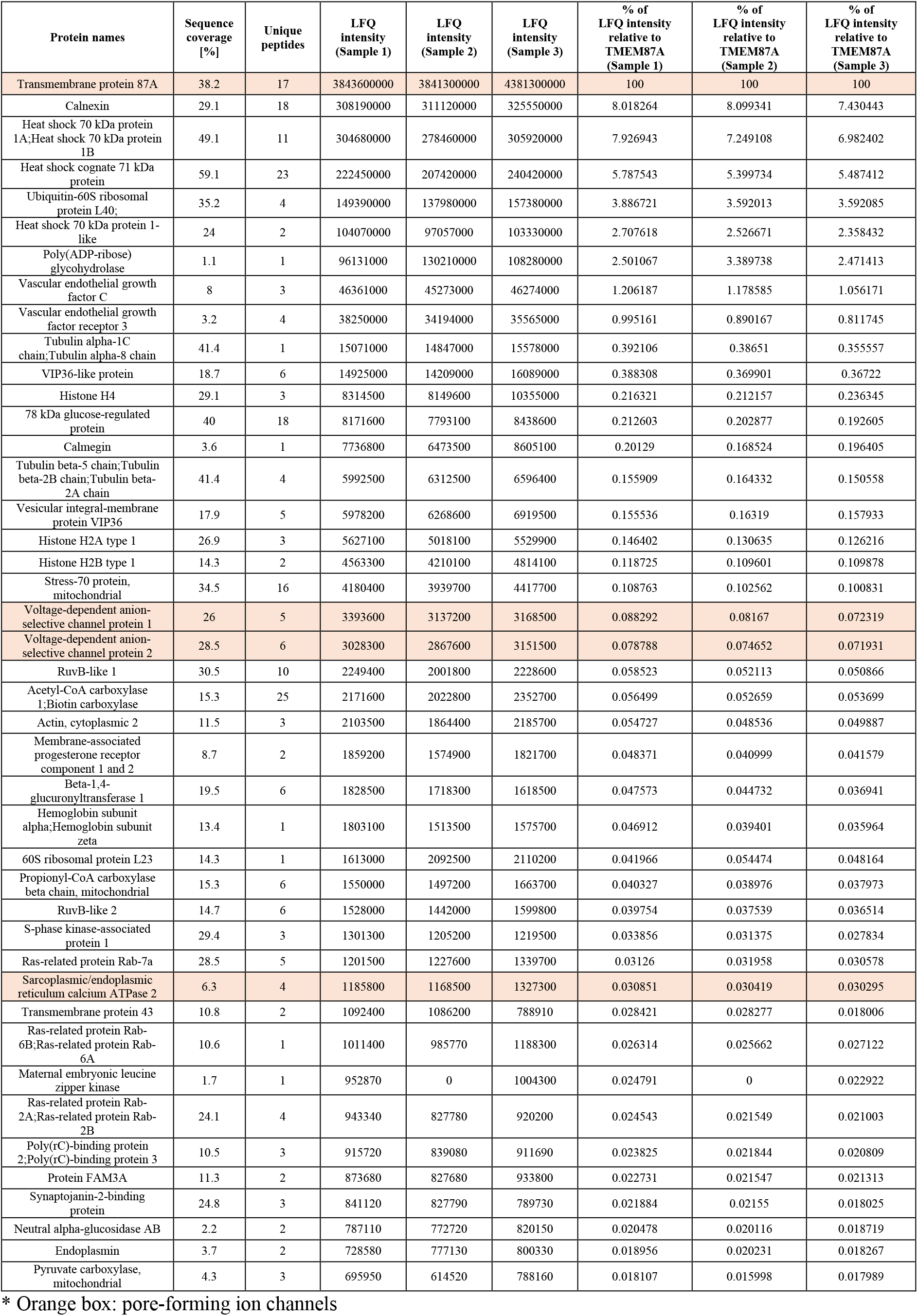
Mass spectrometry of the purified EGFP-tagged TMEM87A solution.

### GolpHCat is required for normal Golgi morphology in hippocampal astrocytes and neurons

To characterize the *in vivo* functions of GolpHCat in mice, we generated a GolpHCat KO mouse line with CRISPR/Cas9 gene editing technique (Fig. 4a). *Tmem87A* of Mus musculus is located on the reverse strand of mouse chromosome 2 (Ch2, 120,185,793-120,234,594) and consists of 20 exons (Fig. 4a). Two guide RNAs (gRNAs) targeting intron 9 and intron 10 to delete the exon 10 containing the GYG motif sequence were designed, and it made a 255 bp deletion in the gene (Fig. 4a). To confirm the generated mouse genotypes, we performed PCR genotyping around the deletion and observed the three types of genotypes: wild-type (WT) with one band of 484 bp, heterozygote (HT) with two bands of 484 bp and 229 bp, and Homozygote (KO) with one band of 229 bp (Fig. 4b). GolpHCat HT and KO mice were viable with no gross abnormality, possibly due to a compensation by TMEM87B.

**Figure 4.**
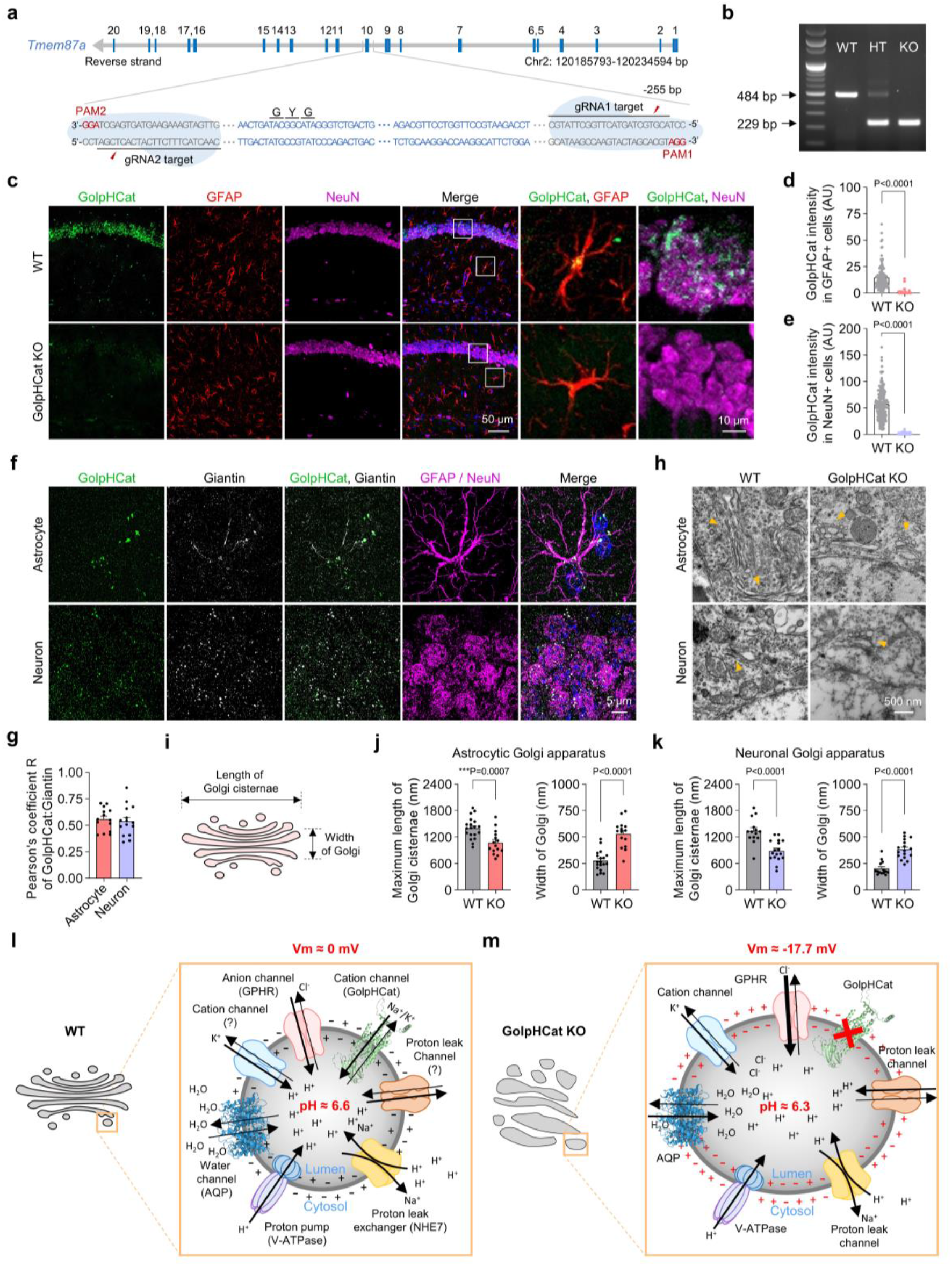
Disruption of Golgi morphology in astrocytes and neurons of GolpHCat KO mice. **a,** Schematic diagram of *Tmem87a* genetic locus on reverse strand of mouse chromosome 2. Exons are indicated with blue box with number, and introns are indicated by gray line (top). Target genomic locus of two gRNAs on intron 9 and 10 are noted with blue, with protospacer adjacent motif (PAM) in the red (bottom). **b,** Representative PCR Genotyping result of wild-type, GolpHCat heterozygote and homozygote mice. **c,** Immunostaining for GolpHCat, GFAP, and NeuN in the hippocampus of WT and GolpHCat KO mice (left). High magnified images showing colocalization of GolpHCat with GFAP+ or NeuN+ cells (right). **d,** Fluorescence intensity of GolpHCat immunoreactivities in GFAP+ cells of WT (n=207 from 4 mice) and GolpHCat KO (n=103 from 3 mice). **e,** Fluorescence intensity of GolpHCat immunoreactivities in NeuN+ cells of WT (n=241 from 4 mice) and GolpHCat KO (n=260 from 3 mice) mice. **f,** Colocalization of GolpHCat with Giantin in hippocampal astrocyte (GFAP) and neuron (NeuN) of WT mice in high-magnification super-resolution microscopy images. **g,** Pearson’s correlation coefficient for colocalization of GolpHCat and Giantin in hippocampal astrocytes (n=14) and neurons (n=15) of WT mice. **h,** TEM images of the Golgi apparatus in hippocampal astrocytes and neurons of WT and GolpHCat KO mice. Yellow arrows indicate Golgi. **i,** Diagram of Golgi structure for analysis used in (**j,k**). **j-k,** Maximum length of Golgi cisternae (left) and width of Golgi (right) in hippocampal astrocytes of WT (n=19 from 3 mice) and GolpHCat KO (n=15 from 3 mice) mice and neurons of WT (n=14 from 3 mice) and GolpHCat KO (n=17 from 3 mice) mice.**l-m,** Hypothetical molecular mechanistic model for Golgi homeostasis in WT (**l**) and GolpHCat KO (**m**) mice. GolpHCat and AQP structures were obtained from AlphaFold structure database. Data are presented as the mean ± SEM. Mann-Whitney test in (**d,e,k-**width). Student two-tailed unpaired t-test in (**j-**length,**j-**width,**k-**length).

To examine the expression pattern of GolpHCat in WT and GolpHCat KGYGO mice, we performed immunohistochemistry (IHC) with antibodies against GolpHCat and GFAP or NeuN, which are astrocytic and neuronal markers respectively, and obtained fluorescent images from hippocampal brain slices (Fig. 4c). The observed GolpHCat immunoreactivities in both GFAP and NeuN positive cells in WT, were virtually absent in GolpHCat KO mice (Fig. 4d, e). These results indicate that the antibody against GolpHCat is highly specific and GolpHCat is majorly expressed in both astrocytes and neurons of the hippocampus.

To investigate whether GolpHCat is localized in Golgi of astrocytes and neurons in a mouse brain, we performed IHC with antibodies against GolpHCat and Giantin, and obtained the super-resolution microscopic fluorescence images (Fig. 4f). We found that GolpHCat was colocalized with Giantin in both cell types (Fig. 4f) with Pearson’s coefficient of 0.56 ± 0.03 for GFAP positive astrocytes and 0.54 ± 0.04 for NeuN positive neurons (Fig. 4g), indicating that GolpHCat is highly localized in Golgi of astrocytes and neurons.

It has been previously reported that genetic ablation of Golgi-pH regulating anion channel, GPHR, disorganized the Golgi structure with fragmentation and swelling of cisternae^10^. To examine the Golgi morphology in hippocampal astrocytes and neurons of WT and GolpHCat KO mice, we performed transmission electron microscopy (TEM) analysis (Fig. 4h). We observed that most of the Golgi exhibited disrupted stacks and dilated cisternae in both cell types of GolpHCat KO compared to WT mice (Fig. 4h). In addition, we analyzed the length of Golgi cisternae and width of Golgi (Fig. 4i) in each cell types and found that the maximum length of cisternae was significantly decreased, whereas width of Golgi was significantly increased in both hippocampal astrocytes and neurons of KO mice (Fig. 4j, k). This is consistent with the morphological alterations observed in GPHR KO cells^10^. In contrast, mitochondria showed normal morphology in KO mice (Extended Data Fig 5a). Taken together, these results indicate that GolpHCat is critical for maintaining the normal Golgi morphology in cells.

Based on these results, we constructed a hypothetical molecular mechanistic model of how Golgi homeostasis is maintained and how GolpHCat contributes to it. The membrane potential of Golgi membrane is extremely difficult to measure, but proposed in previous studies to be 0 mV, +38 mV, or very negative^33,34^. Assuming Golgi membrane potential of 0 mV, GolpHCat functions as a counter cation channel, along with GPHR, the counter anion channel, to maintain the resting pH of 6.6 (Fig. 4l). When GolpHCat is removed, the lack of GolpHCat aberrantly drives excessive Cl^-^ and H^+^ influx into the lumen through GPHR and NHE7, respectively, lowering the resting pH to 6.3 (Fig. 4m). Under this condition, the Golgi membrane potential can be calculated by using Nernst Equation (with the pH shift from 6.6 to 6.3) to be Vm ≈ −17mV (Fig. 4m; see Methods). Simultaneously, the excessive Cl^-^ and H^+^ influx allows water molecules to flow in through the aquaporins (AQPs) due to an osmotic pressure, causing the Golgi to swell (Fig. 4m). Taken together, our mechanistic model predicts that the lack of GolpHCat would result in impaired Golgi homeostasis, leading to altered Golgi functions such as glycosylation and protein trafficking.

### GolpHCat is required for normal glycosylation and protein trafficking

To investigate glycosylation patterns, we analyzed glycans by LC-MS with hippocampal brain samples of WT and GolpHCat KO mice (Fig. 5a), as previously described^35,36^. We found that among the detected 99 N-glycans some were down-regulated, while others were up-regulated in KO mice (Fig. 5a). We performed a multivariant analysis using the partial least squares-discriminant analysis (PLS-DA) and found a complete separation of WT and KO mice (Fig. 5b), indicating that N-glycome was significantly altered in KO mice. To examine the specific change of N-glycome in KO mice, we analyzed relative abundance of each N-glycan type including high-mannose (HM), complex/hybrid (C/H), fucosylated complex/hybrid (C/H-F), sialylated complex/hybrid (C/H-S), and fucosylated/sialylated complex/hybrid (C/H-FS) in WT and KO mice (Fig. 5c). We found that the normalized absolute peak intensity (NAPI) of the most abundant fucosylated complex/hybrid (C/H-F) was slightly and significantly lower in KO mice (Fig. 5c). Furthermore, we analyzed number of fucose in C/H-F and found that NAPI of mono-fucosylated glycan (1) in KO was significantly higher, whereas NAPI of multi-fucosylated glycans (>2) was significantly lower in KO, compared to WT (Fig. 5d). For example, NAPI of mono-fucosylated glycans, 3_4_1_0 and 4_5_1_1, was significantly higher in KO, whereas NAPI of multi-fucosylated glycans, 5_4_3_0 and 6_5_4_0, was significantly lower in KO, compared to WT (Fig. 5e). These results indicate that the fucosylation pattern was selectively and significantly altered in hippocampus of KO mice. To investigate the detailed hippocampal distribution of one of the fucosylated glycans, we performed Matrix-Assisted Laser Desorption Ionization Mass Spectrometry Imaging (MALDI MSI) of tri-fucosylated glycan, 5_4_3_0 (Fig. 5f). We observed that 5_4_3_0 was highly enriched in dentate gyrus (DG) and CA1 of WT hippocampus, which were significantly reduced in the KO (Fig. 5f, g). Taken together, these results indicate that lack of GolpHCat causes altered glycosylation, specifically fucosylation, in the CA1 and DG of the hippocampus.

**Figure 5.**
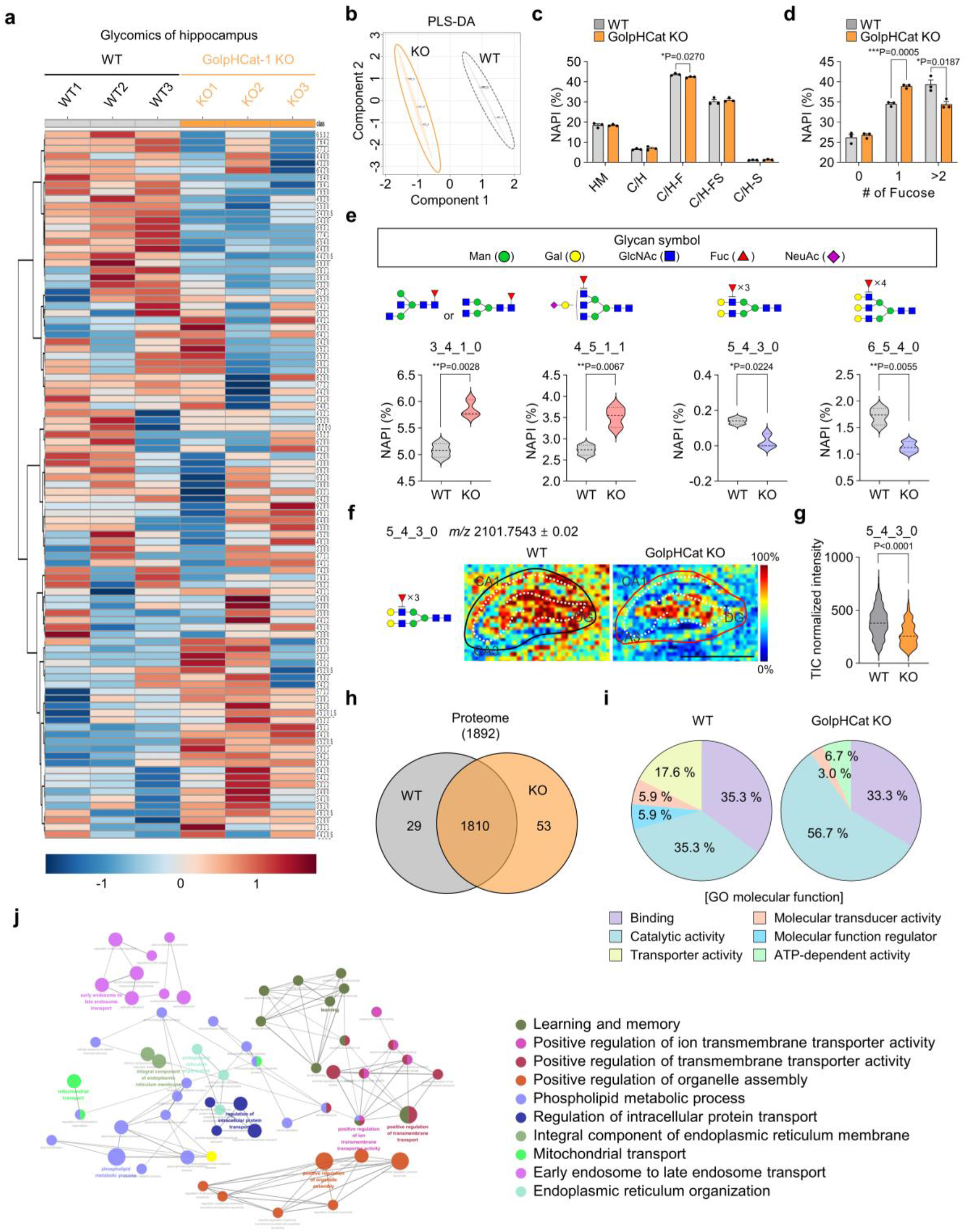
Altered glycosylation and protein trafficking in hippocampus of GolpHCat KO mice. **a-e,** Comparison of 99 N-glycans found in the hippocampus of WT (n=3 mice) and GolpHCat KO (n=3 mice) mice. **a,** Heat map for the hierarchical clustering of N-glycans. Scale bar indicates z-scores of standardized glycan value with higher or lower expressed glycans depicted in red or blue, respectively. (Sugar code: Hex_HexNAc_Fuc_NeuAc_(HexA)_sulfation(S)). **b,** PLS-DA score plots of hippocampus samples between WT and GolpHCat KO. **c,** Glycan biosynthesis types. (high-mannose (HM); complex/hybrid (C/H); fucosylated complex/hybrid (C/H-F); fucosylated and sialylated complex/hybrid (C/H-FS); sialylated complex/hybrid (C/H-S)). **d,** Comparative distribution of all fucosylated glycans NAPI by the number of fucose residues. **e,** Increased and decreased fucosylated glycans in GolpHCat KO compared to WT. (Green circle, Mannose; yellow circle, Galactose; blue square, GlcNAc; red triangle, fucose; diamond magenta, NeuAc). **f,** Spatial distribution of representative N-glycan, 5_4_3_0, from the hippocampus of WT (black line) and GolpHCat KO (red line) imaged by MALDI MSI. White dotted lines indicate the approximate positions of CA1, CA3 and DG. **g,** Total ion count (TIC) normalized intensity from (**f**). **h,** Venn diagram comparison of identified proteins in hippocampus of WT (n=3) versus GolpHCat KO (n=3). **i-j,** Gene Ontology (GO) term analysis with the differentially expressed proteins of WT and GolpHCat KO. **i,** Pie chart depicting the percentage of involvement of identified proteins and their molecular functions. **j,** Network clusters performed with differentially expressed proteins of WT and GolpHCat KO in Cytoscape using ClueGO and CluePedia. The size of the nodes corresponds to the number of the proteins associated to a term. Data are presented as the mean ± SEM. Unpaired t-test in (**c,d,e**).

To investigate protein trafficking patterns, we performed proteomics analysis with hippocampal brain samples of WT and KO mice using LC-MS (Extended Data Fig. 5b). We found 29 differentially expressed proteins present only in WT and 53 differentially expressed proteins only in GolpHCat KO mice (Fig. 5h, Table 2, 3). To examine the role of these differentially expressed proteins, we analyzed molecular function by Gene Ontology (GO) analysis (Fig. 5i). We found that transporter activity was significantly decreased but ATP-dependent and catalytic activity were increased in KO mice (Fig. 5i). To predict the effect of these differentially expressed proteins on biological functions, we performed protein network cluster analysis based on the GO analysis (Fig. 5j). The results showed that these differentially expressed proteins were predicted to have important roles in transporter activity and protein transport of endosome, mitochondria, and endoplasmic reticulum (Fig. 5j). More importantly, these differentially expressed proteins were predicted to play an important role in learning and memory (Fig. 5j). Taken together, these results indicate that the lack of GolpHCat causes altered protein trafficking and glycosylation pattern in hippocampus, and is expected to alter biological functions, especially the hippocampus-related learning and memory.

**Table 2.** Differentially expressed proteins found in WT using mass spectrometry.

| Differentially expressed proteins in WT |
| --- |
| Lysocardiolipin acyltransferase 1 |
| Aladin |
| Envoplakin |
| Rootletin |
| Protocadherin gamma-A4 |
| Sodium-dependent phosphate transporter 2 |
| Long-chain specific acyl-CoA dehydrogenase, mitochondrial |
| Endophilin-B1 |
| Diacylglycerol kinase epsilon |
| Phosphoenolpyruvate carboxykinase [GTP], mitochondrial |
| Signal peptidase complex subunit 2 |
| Metaxin-2 |
| Copine-2 |
| Adenylate kinase isoenzyme 1 |
| Potassium voltage-gated channel subfamily C member 1 |
| Paraplegin |
| Guanylate cyclase D |
| Rho GTPase-activating protein 32 |
| Inward rectifier potassium channel 4 |
| Polypeptide N-acetylgalactosaminyltransferase 16 |
| Carboxypeptidase D |
| A-kinase anchor protein 1, mitochondrial |
| Calcium signal-modulating cyclophilin ligand |
| Kelch repeat and BTB domain-containing protein 11 |
| Glypican-4;Secreted glypican-4;Glypican-6;Secreted glypican-6 |
| Annexin A1 |
| Ig mu chain C region |
| Abelson tyrosine-protein kinase 2 |
| Zinc finger ZZ-type and EF-hand domain-containing protein 1 |

**Table 3.**
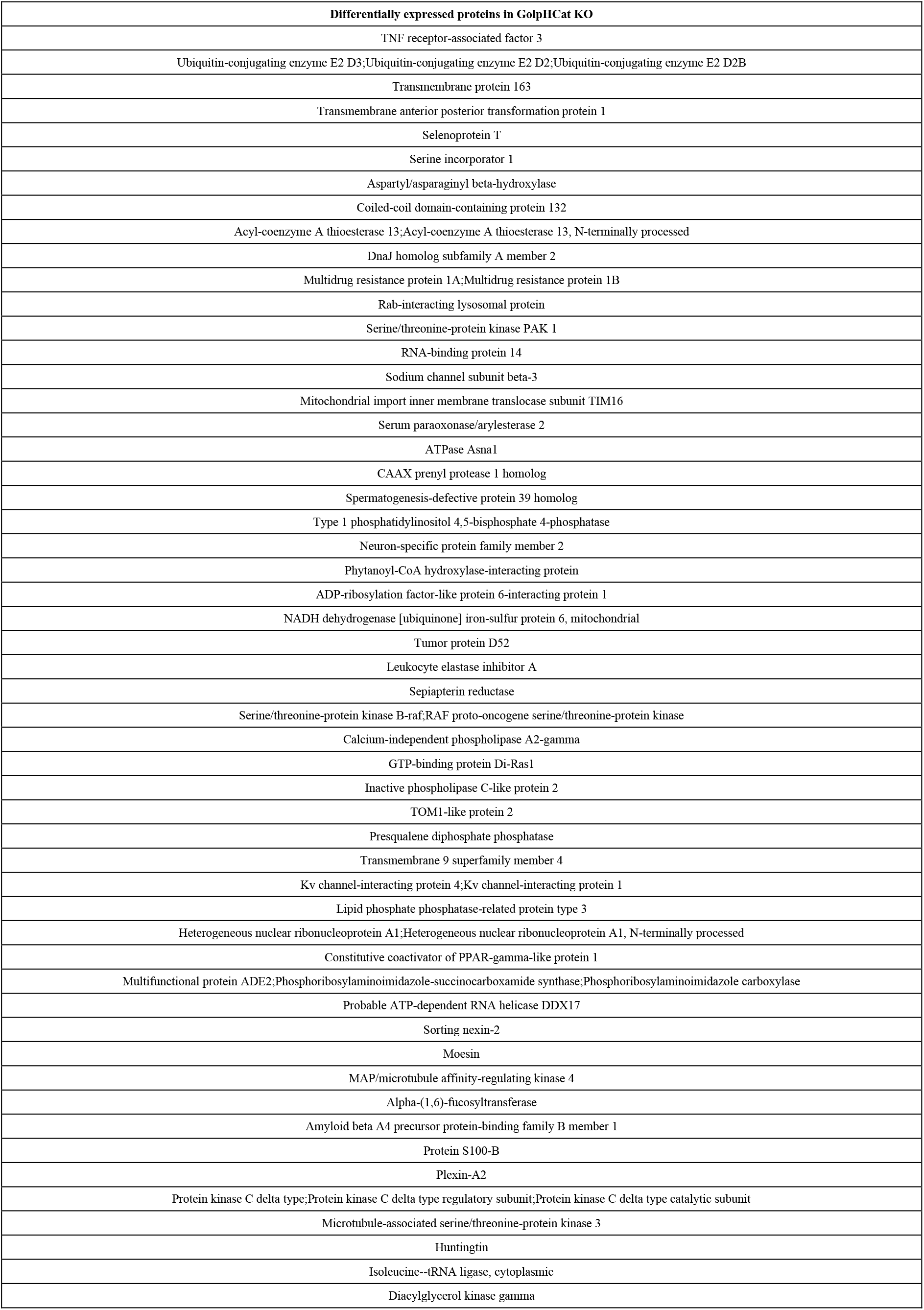
Differentially expressed proteins found in GolpHCat KO using mass spectrometry.

### GolpHCat is required for normal hippocampal memory

To test if the lack of GolpHCat causes altered hippocampus-related learning and memory, we performed detailed electrophysiological and behavioral analyses of the well-established hippocampal spatial memory circuit of CA3→CA1 synapses. Unexpectedly, the intrinsic neuronal excitability of the CA1 pyramidal neurons was not altered in GolpHCat KO compared to WT (Extended Data Fig. 6a-c), although the shape of action potential was slightly altered in KO (Extended Data Fig. 6d-i). To examine the synaptic transmission and plasticity, we performed extracellular field excitatory postsynaptic potentials (fEPSP) at the CA3-CA1 synapses of WT and GolpHCat KO mice (Fig. 6a). We then examined the basal synaptic transmission with increasing stimulus intensity at the Schaffer collateral and again found no difference between WT and GolpHCat KO mice (Fig. 6b). To examine the presynaptic release probability, we measured paired pulse ratios (PPRs) with increasing interpulse intervals and consistently found no significant difference between WT and KO mice at any interpulse intervals (Fig. 6c), These results indicate that the basal synaptic connectivity and transmission were unaltered in KO mice. Finally, to examine the potential role of GolpHCat in synaptic plasticity, we performed high-frequency stimulation (HFS)-induced LTP and found that LTP was significantly impaired in GolpHCat KO mice (Fig. 6d, e). Taken together, these results indicate that the lack of GolpHCat causes altered hippocampal LTP, while leaving the basal synaptic transmission unaltered. At last, to investigate whether the lack of GolpHCat affects hippocampus-dependent memory, we subjected WT and GolpHCat KO mice to hippocampus-dependent spatial memory-related behavioral tasks, such as novel place recognition (NPR) and contextual fear test (Fig. 6f). GolpHCat KO mice showed a significant impairment of both contextual spatial memory in NPR (Fig. 6g, h) and contextual fear memory in fear test (Fig. 6i), with no change of anxiety level in elevated plus maze test (Extended Fig. 7a-f). Taken together, these results indicate that GolpHCat is required for hippocampal spatial and contextual memory, which is consistent with the predictions based on the glycomics and proteomics analyses (Fig. 5f-j).

**Figure 6.**
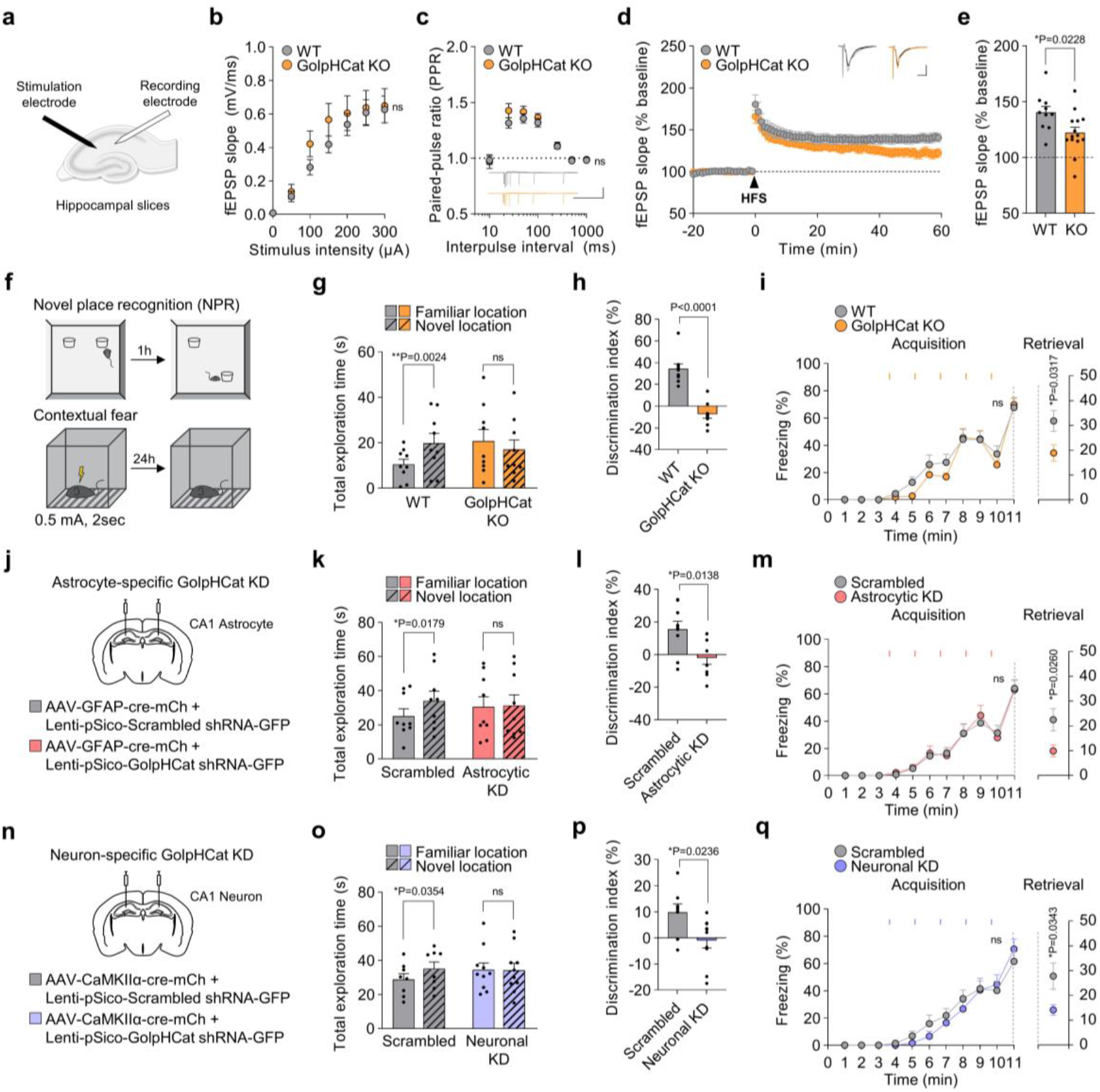
GolpHCat contributes to hippocampal spatial and contextual memories. **a,** Schematic diagram of fEPSP recording in Schaffer-collaterals pathway. **b,** Input-output curve for fEPSP slope obtained with increasing stimulus intensities in WT (n=10 from 3 mice) and GolpHCat KO (n=14 from 3 mice) mice. **c,** Paired pulse ratios (PPR) obtained with increasing interpulse intervals. Inset: representative fEPSP traces. Scale bar: 1 mV and 500 ms. **d,** HFS (1 s at 100 Hz)-induced LTP. Inset: representative fEPSP traces from before and after HFS-induced LTP. Scale bar: 0.5 mV and 10 ms. **e,** Slope of fEPSP over the last 5 min. **f,** Schematic diagram of the novel place recognition (NPR) and contextual fear task. **g,** Total object exploration time during the test phase of NPR task in WT (n=9) and GolpHCat KO (n=9) mice. **h,** Discrimination index during the test phase of NPR task. **i,** Percentage of freezing during acquisition phase of contextual fear task (left) and retrieval phase during the 5 min (right). The orange rectangular indicates the time point of shock. **j,n,** Illustration of virus injection for Scrambled (n=9) and astrocyte-specific GolpHCat KD (Astrocytic KD; n=9) in the stratum radiatum (**j**) or for Scrambled (n=8) and neuron-specific GolpHCat KD (Neuronal KD; n=10) in the pyramidal layer (**n**) of the hippocampal CA1 region. **k,o,** Total object exploration time during the test phase of NPR task. **l,p,** Discrimination index during the test phase of NPR task from (**k,o**). **m,q,** Percentage of freezing during the acquisition phase of contextual fear task (left). Percentage of freezing during the retrieval phase during the 5 min (right). The pink and purple rectangular indicate the moments of shock, respectively. Data are presented as the mean ± SEM. Two-way ANOVA followed by Holm-Šídák’s multiple comparisons test in (**b,c,i-**acquisition**,m-**acquisition**,q-**acquisition), Unpaired t-test with Welch’s correction in (**e,m-**retrieval), Paired t-test in (**g,k,o**), Mann Whitney test in (**h,q-** Retrieval), Unpaired t-test with Welch’s correction in (**i-**retrieval).

Considering the fact that GolpHCat is expressed in both hippocampal astrocytes and neurons (Fig. 4), we investigated which cell type is majorly contributing to the memory impairment in GolpHCat KO mice. We performed cell-type specific gene-silencing (KD) using GolpHCat-shRNA carrying viruses, followed by behavioral tests (Fig. 6j-q and Extended Data Fig. 7a, b). A Cre-dependent shRNA expressing virus (Lenti-pSico-scramble/sh GolpHCat-EGFP) and a cell type-specific Cre-expressing virus (AAV-GFAP-Cre-mCh for astrocyte; Fig. 6j, Extended Data Fig. 7c or AAV-CaMKIIα-Cre-mCh for neuron; Fig. 6n, Extended Data Fig. 7d) were co-injected in hippocampal CA1 region. Both astrocytic and neuronal GolpHCat gene-silencing mice showed impaired spatial and contextual memory in NPR (Fig. 6k, l, o, p) and contextual fear test (Fig. 6m, q). Taken together, these results indicate that GolpHCat in both astrocytes and neurons is critical for hippocampal spatial and contextual memory.

## Discussion

In this study, we have identified an unprecedented unique type of voltage-gated non-selective cation channel, which we named GolpHCat. The single channel analysis of GolpHCat demonstrates that, unlike any other known voltage-gated channels, GolpHCat shows a skewed U-shaped voltage-dependent activation curve, centered around at 0 mV (Fig. 3c). Other voltage-gated channels usually show a unidirectional sigmoidal activation curve^37^, whereas GolpHCat displays a unique bidirectional activation curve at both negative and positive potentials. The one-and-only resembling channel is perhaps the voltage-dependent anion channel (VDAC), which displays, however, an upside-down U-shaped voltage-dependent activation, centered around at 0 mV^38,39^. Based on this unique voltage-activation property, one can make several interesting conjectures about how the Golgi membrane potential is maintained by GolpHCat; 1) GolpHCat’s primary function might be to clamp the resting Golgi membrane potential at 0 mV by opening at both negative and positive offset voltages and resetting the voltage to near 0 mV (by depolarization and hyperpolarization upon channel opening, respectively), 2) the unique U-shaped voltage-dependent activation should render the resting Golgi membrane potential set to 0 mV, independent of the concentration changes in luminal Na^+^ and K^+^ ions, and 3) GolpHCat might be able to achieve this voltage-clamping effect, regardless of the presence of any leak channels.

This important voltage-clamping action of GolpHCat should also be dependent on the luminal pH due to its pH-sensitivity (Fig. 2n, o): the voltage-clamping action of GolpHCat should be maximal at basic luminal pH and minimal at acidic pH as the pH-sensitive domain of GolpHCat should face the lumen (Fig. 4l, m). Therefore, under more basic luminal pH the Golgi membrane potential is likely to be clamped at 0 mV, whereas under more acidic luminal pH it is likely to drift away from 0 mV. Assuming the normal Golgi pH of 6.6 (Fig. 1i), one can further raise a possibility that GolpHCat can serve as a floor to prevent further acidification: when pH is shifted to more acidic pH than the normal Golgi pH, the Golgi membrane potential drifts away to more positive potential due to minimal contribution of GolpHCat’s voltage clamping action, thus allowing proton efflux via the elusive proton leak channel. At the same time, GolpHCat can also prevent further alkalization: at more basic pH, GolpHCat enhances acidification by clamping the Golgi membrane potential to 0 mV and decreasing proton efflux via the elusive proton leak channel (Fig. 4l). Proper measurements of Golgi membrane potential and ion concentrations have been extremely difficult due to the technical challenges of accessing Golgi membrane and structure. Nevertheless, these exciting possibilities should be tested upon imminent technological breakthroughs.

Based on the proposed role of GolpHCat as a unique cation channel that clamps the membrane potential of Golgi to 0 mV, our study enlightens us with the importance of the complex interplay between the membrane potential, ionic balance, and osmotic homeostasis for the maintenance of Golgi structure and volume (Fig. 4l, m). Upon the removal of GolpHCat, the lack of GolpHCat induces disrupted ionic balance with excessive Cl^-^ and H^+^ influx, allowing the excessive flow of water into Golgi lumen due to increasing luminal osmolarity, causing Golgi to swell. Consistent with this proposed mechanism of volume regulation, we observed fragmented and swollen Golgi morphology in GolpHCat KO (Fig. 5h). The swollen transport vesicles might affect delayed and altered protein trafficking. It is consistent with our finding of differentially expressed proteins in either WT or GolpHCat KO that play a role in intracellular protein transport and vesicle transports such as mitochondria, endosome, and ER (Fig. 5h-j). The well-organized ribbon shaped Golgi structure and Golgi pH homeostasis are necessary for proper glycosylation steps in each stack of Golgi, by regulating the exact distribution and activities of glycosylation enzymes, respectively. It is possible that distribution and activities of fucosyl-transferase enzymes may be altered in GolpHCat KO, leading to alteration pattern of fucosylated glycans (Fig. 5a-e). Future work is needed to investigate whether fucosyl-transferase enzymes are altered in GolpHCat KO. Taken together, our results propose that GolpHCat is the key molecule for maintenance of Golgi homeostasis including the Golgi structure and volume, leading to normal Golgi functions such as glycosylation, especially fucosylation, and protein trafficking.

We began our expedition to search for the unidentified counter cation channel in the Golgi. However, what we ended up was an identification of a unique voltage-gated, pH-sensitive, non-selective cation channel, GolpHCat. This still leaves the remaining identification of the counter cation channel. Future studies are needed to identify mysterious yet-to-be discovered ion channels that orchestrate the Golgi homeostasis, which is critical for proper protein glycosylation and trafficking.

## Methods

### Sequence analysis

The partial sequence alignment of the selectivity filter of the potassium channels with hTMEM87A/B and the multiple sequence alignment of TMEM87A isoforms was performed using Clustal Omega (https://www.ebi.ac.uk/Tools/msa/clustalo/). Accession numbers are as follows: hTMEM87A isoform 1, NP_056312.2; isoform 2, NP_001103973.1; isoform 3, NP_001273416.1; hTMEM87B, NP_116213.1; Kv1.1, NP_000208.2; TWIK-1, NP_002236.1; TASK-1, NP_002237.1; TRAAK, NP_201567.1; THIK-1, NP_071337.2; HCN1, NP_066550.2.

Hydrophobicities of human and mouse TMEM87A were calculated in ProtScale (https://web.expasy.org/protscale/) using the Kyte&Doolittle, window size:21, 100%. The signal peptide in the N-terminal of TMEM87A and cleavage site position were predicted in the SignalP-5.0 (https://services.healthtech.dtu.dk/service.php?SignalP-5.0)

### Cell culture

Chinese hamster ovary-K1 (CHO-K1, referred to as CHO) cells and human astrocytes were purchased from the Korean Cell Line Bank and abm (T0280, Richmond, BC, Canada), respectively. CHO cells and human astrocytes were maintained in F-12 (#21127-022, Gibco), and DMEM (#10-013, Corning) respectively. Both media were supplemented with 25g/L glucose, 4g/L L-glutamine, 1g/L sodium pyruvate, 10% heat-inactivated fetal bovine serum (FBS, #10082-147, Gibco), and 1% penicillin-streptomycin (#15140-122, Gibco). Cells were incubated in humidified 5% CO2 incubator at 37 °C.

### Construct and shRNA cloning for *TMEM87A*

Open reading frame of human *TMEM87A* -transcript variant 1 (NM_015497; SC108269), 3 (NM_001286487; SC336449), and mouse *TMEM87A* -transcript variant 1 (NM_173734; MC201598) were purchased from Origene. The coding sequences of these genes were PCR amplified and then subcloned into CMV-pIRES2-DsRed/iRFP vector using the SalI/BamHI restriction enzyme sites or CMV-EGFP-N1 vector using the EcoRI/AgeI restriction enzyme sites using the cloning kit (EZ-Fusion™ HT Cloning core Kit, EZ016TS, Enzynomics). Mutant forms, sh-RNA insensitive form, and isoform 1 truncation of human *TMEM87A* were made using the mutagenesis kit (EZchange™ Site-directed Mutagenesis kit, EZ004S, Enzynomics). Information on oligomer sequences for cloning and mutagenesis were listed in Table 4.

**Table 4.**
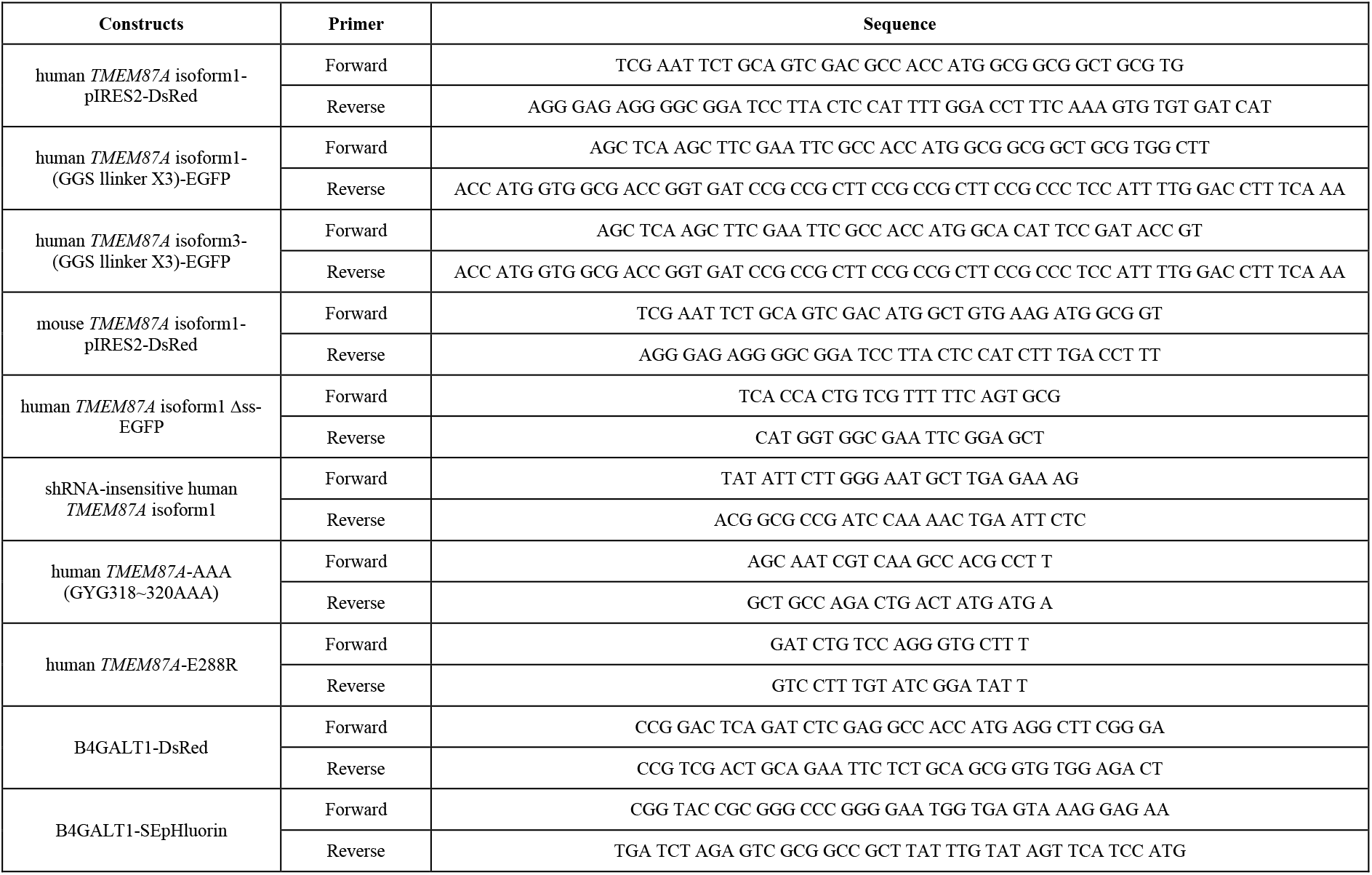
Oligomers used for gene cloning and mutagenesis.

The pSicoR and pSico vectors were used for shRNA knock down in vitro and in vivo, respectively. The targeted sequences for human TMEM87A and mouse TMEM87A were 5’*-GGATTGGTGCTGTCATCTTCC-3’* and *5 ‘-GGATTGGTGCTGTCATCTTTC-3’*, respectively.

For making the human TMEM87A clone construction that is not sensitive to shRNA (shRNA-insensitive human TMEM87A), six nucleotides that make the same amino acid due to the redundancy of genetic code were changed in the human TMEM87A shRNA target site (shRNA-insensitive human TMEM87A sequence 5’-*GGATTGGTGCTGTCATCTTCC*-3’ included 6 mismatches).

### Immunocytochemistry for colocalization and confocal imaging

Human astrocytes were cultured on 0.1 mg/ml poly-D-Lysine (PDL; P6407, Sigma-Aldrich)-coated cover slips in 24 wells for 24 hours. Cells were fixed with 4% paraformaldehyde (PFA) in PBS for 20 min at room temperature before washing three times with PBS and incubating in blocking solution (2% donkey serum (Genetex, GTX27475), 2% goat serum (abcam, ab7481) and 0.03% Triton-X100 (Sigma, 93443) for permeabilization in PBS) at room temperature (RT) for 1 hour. For immunostaining, the samples were incubated overnight at 4°C with primary antibodies: rabbit anti-TMEM87A (1:100, Novus Biologicals, NBP1-90531), mouse anti-Giantin (1:100, abcam, ab37266), chicken anti-GFAP (1:200, Millipore, AB5541) diluted in blocking solution. After incubation overnight, the samples were washed three times with PBS and then incubated for 1hour at RT with secondary antibodies: anti-rabbit IgG Alexa Fluor 488 (1:500, Jackson lab), anti-mouse IgG Alexa Fluor 594 (1:500, Jackson lab), and anti-chicken IgG Alexa Fluor 647 (1:500, Jackson lab) diluted in the blocking solution. After secondary antibody incubation, samples were washed three times with PBS and DAPI solution (1:2,000, Pierce, 62248) was added during the second washing step. Finally, samples were mounted with fluorescent mounting medium (Dako, S3023). Fluorescence images were taken using a Zeiss LSM 900 confocal microscope with a 63X lens.

### Protein localization with TMEM87A isoforms and confocal images

Human astrocytes were cultured on PDL-coated cover slips for 24 hours before transfection. Cells were transfected with TMEM87A shRNA-mCh with each shRNA insensitive EGFP-tagged TMEM87A isoform for 48 hours. Cells were washed once with PBS, then fixed with 4% paraformaldehyde in PBS for 20 min at RT and washed three times. DAPI solution was added during the second washing step. Finally, samples were mounted with fluorescent mounting medium. Fluorescence images were taken at 405 nm for DAPI, 488nm for EGFP, and 594nm for mCh using a Zeiss LSM 900 confocal microscope with a 63X lens.

### Illumina Hiseq library preparation and sequencing

RNA was isolated from cultures human astrocytes using Qiagen RNEasy Kit (Cat. No #74106). Sample libraries were prepared using the Ultra RNA Library Prepkit (NEBNext, E7530), Multiplex Oligos for Illumina (NEBNext, E7335) and polyA mRNA magnetic isolation module (Invitrogen, Cat. No. #61011). Full details of the library preparation and sequencing protocol are provided on the website and previously described^40^. The Agilent Bioanalyser and associated High Sensitivity DNA Kit (Agilent Technologies) were used to determine the quality, concentration, and average fragment length of the libraries. The sample libraries were prepared for sequencing according to the HiSeq Reagent Kit Preparation Guide (Illumina, San Diego, CA, USA). Briefly, the libraries were combined and diluted to 2nM, denatured using 0.1N NaOH, diluted to 20pM by addition of Illumina HT1 buffer and loaded into the machine along with read 1, read 2 and index sequencing primers. After the 2×100 bp (225 cycles) Illumina HiSeq paired-end sequencing run, the data were base called and reads with the same index barcode were collected and assigned to the corresponding sample on the instrument, which generated FASTQ files for analysis.

### NGS Data Analysis

BCL files obtained from Illumina HiSeq2500 were converted to FastQ and demultiplexed based on the index primer sequences. The data was imported to Partek Genomics Suite (Flow ver 10.0.21.0328; copyright 2009, Partek, St Louis, MO, USA), where the reads were further processed. Read quality was checked for the samples using FastQC. High quality reads were aligned to the *Homo sapiens* (human) genome assembly GRCh37 (hg19, NCBI using STAR (2.7.8a). Aligned reads were quantified to the human genome assembly transcript model (hg19 - RefSeq transcripts 93) and normalized to obtain fragments per kilobase million (or FPKM) values of positively detected and quantified genes. Alternate splice variants for the genes were detected during quantification and also normalized to obtain FPKM values for the alternatively spliced variants.

### pH measurement with sensor imaging

For the pH measurement, pSicoR TMEM87A shRNA-DsRed and B4GALT1-SEpHluorin were transiently co-transfected into human astrocytes one day before imaging. For the measurement of the Golgi pH buffer capacity, we obtained basal pH images for 5 min and then treated 100 mM NH4Cl for 15 min. Fluorescence live images were excited at wavelengths of 405 nm and the emitted fluorescence was captured through a spectral slit for wavelengths of 470nm and 520nm. The data were imported into Microsoft Excel for calculation of the intensity ratios (520 nm:470 nm) and further analyses. Golgi pH values were estimated from intensity ratios using the calibration curve^41^.

### Electrophysiological recording of TMEM87A

For whole-cell patch clamp recording, TMEM87A WT or mutant-pIRES2-DsRed was transiently transfected into CHO-K1 cells one day before patch recording. Transfection reagent (Lipofectamine 3000; Invitrogen, L3000001) was used for transfection in all experiments. After 24 hours, cells were seeded onto 0.1 mg/ml PDL-coated cover slips and used for whole-cell patch-clamp recording within 12 hours.

Unless otherwise indicated, the bath solution contained (in mM) 150 NaCl, 3 KCl, 10 HEPES, 5.5 glucose, 2 MgCl_2_, 2 CaCl_2_ with pH adjusted to 7.3 by NaOH (320~325 mOsmol/kg). Borosilicate glass pipettes (Warner Instrument Corp., USA, GC150F-10) were pulled and had a resistance of 4–6 MΩ in the bath solution when filled with a pipette solution contained (in mM) 130 K-gluconate, 10 KCl, 10 HEPES, 10 1,2-Bis(2-aminophenoxy)ethane-*N,N,N′,N′*-tetraacetic acid (BAPTA) with pH adjusted to 7.3 by KOH (290~310 mOsmol/kg). Cells were held at −60mV For recording with voltage-clamp ramp protocol, currents were measured under the 1000 ms-duration voltage ramps descending from +100 mV to −150 mV with 10 sec time intervals. For recording with voltage-clamp step protocol, currents were measured under the step pulses from +100mV to - 150mV with 100ms with 1sec time intervals. Electrical signals were amplified using MultiClamp 700B (Molecular Devices, USA). Data were acquired by Digitizer 1550B (Molecular Devices, USA) and pClamp 11 software (Molecular Devices, USA) and filtered at 2 kHz. These machines were used in all electrophysiology experiments in this paper.

To determine the contribution of Na^+^ in the bath solution for the inward currents, 150 NaCl containing bath solution was replaced to 150 NMDG-Cl under the same other condition. For measuring the cation permeability through TMEM87A, 150 Na^+^ in the bath solution was substituted to 150 X, where X was K^+^ or Cs^+^ (pH 7.3 was adjusted with KOH and CsOH, respectively). The pipette solution was prepared as described above. For calculating the relative permeability ratio of TMEM87A, we measured the differences of reversal potential between two cationic I-V curves. The permeability ratio, P_X_ /P_Na_, was estimated using the modified Goldman – Hodgkin - Katz equation as reported previously:

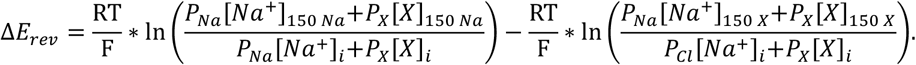

To determine the impermeability of anion, 150 NaCl containing bath solution was replaced to 150 Na-isethionate under the same other condition. The I-V curves in this experiment were corrected with the LJP (LJP: NaCl, +13mV; Na-isethionate, −11.6mV).

To assess the inhibitors as a TMEM87A blocker, they were contained in the bath solution and then recorded using the following concentration: Gadolinium chloride (GdCl_3_) (0.1, 0.3, 1, 3, and 10 μM, Sigma, G7532), Na-gluconate (gluconate) (0.01, 0.03, 0.1, 1, and 10 μM, Sigma, S2064), and ZD7288 (100 μM, TOCRIS, 1000). The IC_50_ values was estimated by non-linear regression analysis, using the following equation:

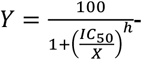
where *X* is the inhibitor concentration, *IC*_50_ is the concentration required for half-maximal inhibition, and *h* is the Hill slope constant (h: gadolinium, −1.498; gluconate, −1.301).

To measure the extracellular pH-dependency of TMEM87A currents, different pH solutions were made using the composition previously described and the pH was manipulated with HCl or NaOH.

### Surface biotinylation assay and western blot

For the surface biotinylation assay, TMEM87A WT-pIRES2-DsRed or TMEM87A-AAA-pIRES2-DsRed was transiently transfected into HEK293T cells on 60mm dishes one day before the experiment. After 24 hours, cells were washed three times with ice cold PBS-CM (PBS-CaCl2/MgCl2) and incubated with 0.5 mg/ml Ez-Link Sulfo-NHS-LC-Biotin solution (Thermo Scientific, 21335) diluted in PBS-CM on the shaking for 30 min at 4°C. After biotinylation, quenching buffer (100mM glycine in PBS-CM) was added to stop the reaction and cells were washed three times with the buffer. Cells were lysed in 250 mL RIPA lysis buffer and incubated 10 min on ice, followed by centrifugation at 14,000 rpm for 10 min at 4°C. The supernatant was collected, and protein concentration was measured using the BCA Protein Assay Kit (Thermo Scienrific, 23225). Volume containing equal protein amounts of each sample were incubated with high capacity Neutravidin Agarose Resin (Thermo Scientific, 29204) overnight at 4°C. The resin was washed three times with lysis buffer and bound proteins were subjected to western blotting. Cell lysates and Biotinylated samples were loaded on 8% SDS-polyacrylamide gels and transferred to PVDF membranes. The membranes were then blocked with 5% skim milk in TBST (20 mM Tris-HCl (pH 7.5), 150 mM NaCl, 0.05% Tween 20) for 1 hour at RT, and then incubated with primary antibodies: rabbit anti-TMEM87A (1:1000, Novus Biologicals, NBP1-90531), and rabbit anti-β-actin (1:2000, Abcam, ab133626) diluted in 5% skim milk in TBST overnight at 4 °C. The next day, membranes were incubated with the corresponding horseradish peroxidase-conjugated secondary antibodies (1:5000, KPL, USA) diluted in 2.5% skim milk in TBST for 1 hour at RT. Immuno-reactive protein bands were detected using ECL Western Blotting Substrate (Thermo Fisher Scientific, USA).

### Cloning, Expression, and purification for liposome patch of hTMEM87A

Human TMEM87A (hTMEM87A, NP_056312.2, M1-E555) followed by TEV protease cleavage sequence (ENLYFQG), PreScission Protease cleavage sequence (LEVLFQGP), EGFP (M1-239K), thrombin cleavage sequence (LVPRGS) and Twin-strep-tag were cloned into the BamHI and XhoI sites of a pcDNA3.4 (#A14697, Invitrogen).

Recombinant hTMEM87A proteins (WT and mutants) were transiently expressed in Expi293F cells (#A14527, Thermo Fisher Scientific) according to the manufacturer’s instructions. Briefly, 200μg of plasmid DNA was transfected into 200 ml of Expi293F cells (3.0 x 106 cells/ml) using Expifectamine (#A14524, Thermo Fisher Scientific). Cells were cultured in Expi293 expression medium (#A14351, Thermo Fisher Scientific) at 37°C and 8% CO2 with shaking (orbital shaker, 120 rpm). After 20 h, the culture was supplemented with enhancer (#A14524, Thermo Fisher Scientific) and further incubated for 30-34 hours at 30°C. Cell pellets were resuspended in 20 ml HN buffer [50mM HEPES pH 7.5, 250mM NaCl, and 1x complete protease inhibitor cocktail (#11836170001, Roche)] and lysed by sonication (total 2min, 1sec with intervals of 5sec, 20% amplitude). After ultracentrifugation (Beckman Ti70 rotor, 150,000 x g for 1 h), the collected membrane fraction was homogenized with a glass Dounce homogenizer in 20ml buffer [HN buffer + 1% (w/v) n-dodecyl β-D-maltoside (DDM; #D310S, Anatrace) and 0.2% (w/v) cholesteryl hemisuccinate (CHS; #CH210, Anatrace)] and solubilized for 2h at 4°C. The insoluble cell debris was removed by ultracentrifugation (Beckman Ti70 rotor, 150,000 x g for 1 h), and the supernatant was incubated with 2ml Strep-Tactin resin (#2-1201-025, IBA Lifesciences) for 30 min at 4°C. After washing with 10 column-volumes of wash buffer [HN buffer + 0.05% (w/v) DDM and 0.01% (w/v) CHS], the hTMEM87A-EGFP-Twin strep tag was eluted with 5ml elution buffer [HN buffer+ 0.05% (w/v) DDM/CHS + 10mM desthiobiotin]. After concentration using Amicon Ultra centrifugal filter (100-kDa cut-off; Millipore), the hTMEM87A-EGFP-Twin strep was further purified by size exclusion chromatography (SEC) using a Superose 6 Increase 10/300 GL column (#29-0915-96, Cytiva) equilibrated with a final buffer [HN buffer + 0.01% (w/v) DDM and 0.002% (w/v) CHS]. The peak fractions were immediately used for preparing proteo-liposomes.

### Reconstitution of human TMEM87A

A total of 10mg of lipids (8:2, POPC : POPG; Avanti Polar Lipids) was dissolved in 1mL chloroform in the glass tube, dried to a thin film under a nitrogen stream, and further dried overnight under a vacuum. A total of 2 mL dehydration/rehydration (D/R) buffer (5 mM Hepes, 200 mM KCl pH 7.2 adjusted by KOH) was added to the lipids and the solution was vortexed for 60sec before being bath sonicated until transparent for 20min. Purified hTMEM87A was added to 2mg lipids in D/R buffer and reconstituted into liposomes at 1:100 protein-to-lipid ratio. D/R buffer was used to bring the volume to 1mL and roller mix at room temperature for 1h. After the beads settled to the bottom of the tube to eliminate detergents, the supernatant was collected and ultracentrifuged for 45 min at 4°C and 250,000g. Pelleted proteoliposomes were resuspended in 80 μL D/R buffer by gently pipetting and used on the day. 3-4 spots of 20 μL of proteoliposomes were placed on a glass coverslip coated with 0.1 mg/ml PDL. Proteoliposomes were vacuum dried at room temperature for 6 h at 4 °C and then rehydrated with 20 μL DR buffer to each spot with wet filter paper overnight at 4 °C (8–24 h) for patch-clamp recording.

### Single-Channel Recording and Analysis

A previously reported protocol (ref) was used for the TMEM87A single channel recordings in the liposome. All recordings were performed with attached liposome patch. For the TMEM87A single channel recording, symmetric solution was used in bath and pipette solutions followed by: (in mM) 200 KCl, 5 HEPES with pH adjusted to 7.2 by KOH. Notably, for making the blister, 40 mM MgCl2 was added in bath solution 30 min before patch and was maintained thoughout the recording. Borosilicate glass pipettes were pulled and polished to a resistance of 3–6 MΩ in the bath solution. The currents were recorded at RT and holding current was indicated in the data. The single channel analysis was performed in Clampfit 10.7.

### Generation of TMEM87A knockout mice

We requested the generation of TMEM87A knockout mice line fromGHBio (Daejeon, Korea). TMEM87A KO allele was generated by using the CRISPR/Cas9-mediated homologous recombination. Two guide RNAs (gRNAs), which bound to intron 9 or intron 10 and deleted the whole exon 10, including the GYG sequence, were designed with the following spacer sequences: spacer of gRNA1, 5’-TAAGCCAAGTACTAGCACGT-3’; spacer of gRNA2: 5’-GATGAAAGAAGTAGTGAGCT-3’. Protospacer adjacent motif (PAM), AGG, for corresponding gRNA was located on the 3 bp downstream of each targeted DNA sequences. The targeting construct was injected into a C57BL/6J mouse ES cell line with the two gRNAs. PCR and sequencing analysis were used to identify ES cell clones with proper targeting. Obtained female knockout mice were maintained by crossing with male wild-type mice. Genotypes were determined by PCR using the following primers; Forward: 5’-TGT GCA CAT AAC TGA GGT CAT-3’; Reverse: 5’-GCT TCA CTG CAA TCT TTC TGC-3’.

### Animals

All mice were maintained under 12:12-h light-dark cycle (lights on at 8:00 AM) and had ad libitum access to food and water. Animal use and procedures were approved by Institutional Animal Care and Use Committees (IACUCs) of IBS (Daejeon, Korea). IHC, behavioral tests, and slice patch were performed using 9- to 15-week male mice and wild-type littermates were used. Virus injection was done using 6- to 7-week male mice.

### Immunohistochemistry and confocal imaging

Adult mice were deeply anesthetized with isoflurane and transcardially perfused with saline, followed by cold 4% paraformaldehyde in 0.1M PBS. Brains were post-fixed in 4% paraformaldehyde for 24 hours at 4°C and 30% sucrose for 48 hours at 4°C. Frozen brains in OCT embedding compound solution were cut into 30 μm coronal sections. Sectioned brains were washed three times in PBS and incubated in blocking solution (2% donkey serum, 2% goat serum and 0.3% Triton-X100 for permeabilization in PBS) at room temperature (RT) for 1 hour. Samples were incubated in the blocking solution (2% donkey serum, 2% goat serum and 0.3% Triton-X100 for permeabilization in PBS) at RT for 1 hour. For immunostaining, the samples were incubated overnight at 4°C with primary antibodies: rabbit anti-TMEM87A (1:100, Novus Biologicals, NBP1-90531), mouse anti-Giantin (1:100, abcam, ab37266), chicken anti-GFAP (1:200, Millipore, AB5541), guineapig anti-NeuN (1:200, Millipore, ABN90) diluted in blocking solution. After incubation overnight, the samples were washed three times with PBS and then incubated for 1hour at RT with secondary antibodies: anti-rabbit IgG Alexa Fluor 488 (1:500, Jackson lab), anti-mouse/chicken IgG Alexa Fluor 594 (1:500, Jackson lab), and anti-chicken/guineapig IgG Alexa Fluor 647 (1:500, Jackson lab) diluted in in blocking solution. After secondary antibody incubation, samples were washed three times with PBS and DAPI solution was added during the second washing step. Finally, samples were mounted with fluorescent mounting medium. Fluorescence images were taken using a Zeiss LSM900 confocal microscope with a 20X and 63X lens and obtained Z-stack images in 1~2μm steps and the fluorescent intensity were analyzed using the Image J software. For colocalization analysis, fluorescence images were taken using the Zeiss Elyra 7 Lattice SIM with 63X lens and Pearson’s coefficient R were analyzed using the colocalization tools in ZEN Blue software.

### Transmission electron microscopy and analysis for Golgi morphology

Brains were fixed with 2% glutaraldehyde and 2% PFA in 0.1M PBS for 12 hours at 4°C and washed in 0.1M PBS, and then post-fixed with 1% OsO4 in 0.1M PBS for 1.5 hours. The samples were then dehydrated with increasing concentrations of ethanol (50-100%),infiltrated with propylene oxide for 10 min and embedded with a Poly/Bed 812 kit (Polysciences, USA) and polymerized for 18 hours at 60°C. The samples were sectioned into 200nm with a diamond knife in the ultramicrotome (EM-UCT, Leica, USA) and stained with toluidine blue for observation with optical microscope. Thin sections (70nm) were double-stained with 5% uranyl acetate for 10 min and 1% lead citrate for 5 min. Images were taken using the transmission electron microscope (JEM-1011, JEOL, Japan) at the acceleration voltage of 80kV and photographed with a digital CCD camera (Megaview III, Germany). Golgi morphologies were analyzed using the ZEN Blue software.

### Calculation of predicted Golgi membrane potential of GolpHCat KO

The Golgi membrane potential in the lack of GolpHCat, was estimated using the modified Nernst Equation as reported previously:

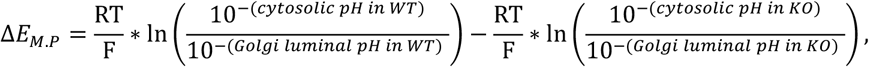

with Golgi pH shift from 6.6 in WT to 6.3 in GolpHCat KO, which are obtained from Fig. 1i. The cytosolic pH is 7.3 in both conditions.

### Slice patch recording for hippocampal neurons and astrocytes

To test intrinsic properties of the hippocampal neurons and astrocytes, slice recording was performed using a modified protocol from the previous reports^42^. Briefly, brain was excised from the skull and sectioned in an ice-cold, oxygenated (95% O_2_/5% CO2) sucrose-based dissection buffer containing (in mM) 212.5 sucrose, 5 KCl, 1.23 NaH2PO4, 26 NaHCO3, 10 glucose, 0.5 CaCl2, 10 MgSO4, pH 7.4. Brain slices were transversely cut into 300 μm thick sections containing hippocampus using vibrating microtome (DSK Linearslicer™ Pro7, DSK, Japan). Prepared brain slices were recovered and recorded in oxygenated (95% O2/5% CO2) artificial cerebrospinal fluid (aCSF) containing (in mM) 124 NaCl, 5 KCl, 1.25 NaH2PO, 26 NaHCO3, 2.5 CaCl2, 1.5 MgCl2 and 10 glucose for at least 1 hour at 28±1° prior to recording. Brain slices for astrocyte patch were co-loaded with 0.5 μM SR-101 (Sigma; S7635) dye to identify the location of astrocytes in the CA1 region.

For rheobase and action potential recording in hippocampal pyramidal neurons, patch electrode (6~8 MΩ) was filled with an internal solution (in mM): 145 K-gluconate, 10 HEPES, 5 KCl, 0.2 EGTA, 5 Mg-ATP, and 0.5 Na_2_-GTP,pH adjusted to 7.3, and osmolarity 295 mOsmol/kg). Measurement was performed in a whole-cell current-clamp configuration, with no membrane potential adjustment. Rheobase and action potential were measured by giving 5 pA or 20 pA depolarizing steps for 1 s injection with 3 s between steps, respectively. Frequency, spike half-width, spike rise, and spike decay values of of Rheobase were analyzed by Minianalysis software (Synaptosoft). For passive conductance recording in hippocampal astrocytes, measurement was performed in a whole-cell voltage-clamp configuration and we used the holding potential of −80mV. Patch electrode (6~8 MΩ) was filled with an internal solution (in mM): 140 KCl, 10 HEPES, 5 EGTA, 2 Mg-ATP, 0.2 NaGTP, adjusted to pH 7.4 with KOH). Currents were measured under the 1000 ms-duration voltage ramps descending from +100 mV to −150 mV with 10 s time intervals. Passive conductance data analysis was performed using Clampfit (Molecular Devices)

### Field excitatory postsynaptic potential (fEPSP) recording

To test basal synaptic transmission, paired-pulse ratio (PPR), and long-term potentiation (LTP), brain slice preparation and fEPSP experiments were performed as described previously^42^. Briefly, mouse was anaesthetized with isoflurane and decapitated. Isolated brain from decapitation was cut into 400-μm-thick transverse hippocampal slices using vibrating microtome (DSK Linearslicer™ Pro7, DSK, Japan) in ice-cold, oxygenated (95% O2/5% CO2) sucrose-based dissection buffer containing 5 KCl, 1.23 NaH_2_PO_4_, 26 NaHCO_3_, 10 glucose, 0.5 CaCl_2_, 10 MgSO_4_, and 212.5 sucrose (in mM).

Brain slices were recovered in oxygenated aCSF containing 124 NaCl, 5 KCl, 1.25 NaH2PO4, 2.5 CaCl2, 1.5 MgCl2, 26 NaHCO3 and 10 glucose (in mM) at 28 ± 1°C for at least 1 hour and subjected to the fEPSP recordings. To evoke fEPSP from Schaffer collateral pathway, electrical stimulus was delivered with a concentric bipolar electrode (CBBPE75, FHC, Bowdoin, ME, USA).

To record fEPSP in Schaffer collateral pathway, aCSF-filled recording pipette, fabricated from a borosilicate glass capillary (1–3 MΩ, GC100T-10, Harvard Apparatus, USA), was placed in stratum radiatum of hippocampal CA1. The slope of fEPSP was acquired and analyzed with WinLTP v2.01 software (WinLTP Ltd., The University of Bristol, UK). For the basal synaptic transmission, stimulus intensity was increased by 50 pA from 0 to 300 pA. In subsequent experiments, the stimulus intensity was set to 40-45% of the maximum response. For the PPR experiment, two pulses were delivered at intervals of 10, 25, 50, 100, 250, 500, and 1000 ms, and the ratio was calculated by dividing the fEPSP slope from second response by the one from first response. For the LTP experiment, the slope of fEPSPs was monitored at 0.067Hz (one pulse per 15 seconds) during the experiment. After obtaining a stable fEPSP response for at least 20 minutes, LTP was induced with a single high-frequency stimulation (HFS) (1 s at 100 Hz). To quantify the degree of potentiation, the fEPSP slopes over the last 5 min from each slice were averaged.

### Behaviors

#### Novel place recognition behavioral task

Novel place recognition task was performed as previously described^43^. For the experiment, mice were placed in an open field with two identical objects and given 10 minutes to explore these objects and returned to their home cage. After 1 hour, mice were placed back into the open field with two same identical objects with one object relocated to a novel place in the field (corner directly opposite to object’s previous location).Mice were recorded while exposed to this condition for 10 minutes to observe their spatial recognition memory. The Discrimination index was calculated as the percentage of time spent examining the object in the novel place over the total time spent examining both objects.

#### Contextual fear behavioral task

On the training day, mice were placed in a standard fear conditioning shock chamber. Mice were allowed to explore the chamber freely for 5 minutes and then received the first electrical foot shock (0.5mA, 2 seconds duration, followed by four more shocks at 1 minute and 30 seconds intervals. After 24 hours, on the test day, mice were placed back into the same chamber for 5 minutes to assess freezing during the retrieval phase. The mice’s movement in the fear conditioning chamber were recorded using a near-infrared camera and analyzed in real time with EthoVision XT 11 software (Noldus). Freezing behavior was defined as immobility for more than 2 seconds. Freezing percentage was calculated as the immobile time divided by total time.

#### Stereotaxic virus injection into the hippocampal CA1

All viruses used in this study were produced at the Institute for Basic science virus facility (IBS virus facility, Daejeon). Mice were placed in stereotaxic frames after being anesthetized with vaporized isoflurane (Kopf). The scalp was incised, and a hole was drilled into the skull above the CA1 (anterior/posterior, −1.5mm; medial/lateral, ±1.5mm from bregma, dorsal/ventral, −1.8~2.0mm from the brain surface). The virus was loaded into a glass needle and injected bilaterally into the CA1 at a rate of 0.1 μl/min for 5 min (total 0.5μl) using a syringe pump (KD Scientific). In each experiment, AAV-GFAP-Cre-mCh, AAV-CaMKIIa-Cre-mch, Lenti-psico-Scramble-GFP, and Lenti-pSico-TMEM87A shRNA-GFP viruses were used. Three weeks after the virus injection, mice were used for behavioral experiments and Immunohistochemistry.

#### Extraction of membrane protein from brain tissue samples

Brain tissue samples (WT, n=3; GolpHCat KO, n=3) were homogenized with a buffer consisting of 0.25 M sucrose, 20 mM HEPES-KOH pH 7.4, and 1:100 protease inhibitor mixture by sonication. The protein concentration of the homogenized samples was determined using a Qubit 2.0 Fluorometer, and 250 μg of protein was used for membrane extraction. The lysates containing 250 μg protein were pelleted by ultracentrifugation at 200,000 g for 45min in a homogenization buffer, then resuspended in 0.2 M Na2CO3 (pH 11), and centrifugated at 200,000 g for 45min. Finally, the membrane fraction was extracted by once more resuspending 0.2 M Na2CO3 and centrifuging at 200,000 g for 45min^44^.

#### Enzymatic release and enrichment of N-glycans

Each membrane fraction resolubilized in 50 μL deionized (DI) water was mixed with an equal volume of 200 mM NH4HCO3 and 10 mM dithiothreitol (Sigma-Aldrich), and then denatured for thermal cycle (100 °C, 2 min). After samples cooling, 2 μL of peptide N-glycosidase F (New England Biolabs) were added, and entire mixture were incubated in a water bath at 37 °C for 16 h. The mixture was chilled in 80% (v/v) ethanol (Merck) at – 42 °C for 1h, and a glycan-rich supernatant was collected by centrifugating and precipitating out the deglycosylated proteins. The supernatant fraction was dried in the condition of vacuum and followed as purifying the released N-glycans using porous graphitized carbon-solid phase extraction (PGC-SPE). Briefly, PGC cartridges (Agilent Technologies, USA) were first washed with 6 mL of DI water and 6 mL of 80% acetonitrile (ACN) and 0.1% trifluoroacetic acid (TFA) in DI water (v/v). And the cartridge was conditioned with 6 mL of DI water, followed by loading aqueous N-glycan solution onto the cartridge. Continuously, N-glycans were eluted stepwise with 6 mL of 10% ACN (v/v), 20% ACN (v/v), and 40% ACN and 0.05% TFA (v/v) in DI water after washing with 8 mL of DI water. All fractions were dried under vacuum and resolubilized with 15 μL of DI water before LC/MS analysis.

#### N-glycan profiling by nano-LC/Q-TOF MS

Purified glycans were analyzed at nano-LC/Q-TOF MS with condition of a nano-LC chip consisting of porous graphitized carbon analytical column (5 μm, 0.075 × 43 mm i.d.). Glycans were separated at 0.3 μL /min with a 65 min gradient using Buffer A (3.0% ACN with 0.1% formic acid (v/v) in water and Buffer B (90% ACN with 0.1% formic acid (v/v) in water). The LC gradient used was as follow: 2.5 min, 0% B; 20 min, 16% B; 30 min, 44% B; 35 min, 100% B; 45 min, 100% B; 45.01 min, 0% B. MS spectra were acquired in positive ionization mode with a mass range of *m/z* 500-2000 and an acquisition time of 0.63 s per spectrum.

After data acquisition, raw LC/MS data was processed by the Molecular Feature Extractor algorithm of MassHunter Qualitative Analysis software B.07.00 (Agilent Technologies). A list of all N-glycans was extracted using previously optimized application of spatial mouse brain glycome database^35^. N-glycan compositions were identified with a mass error tolerance of 10 ppm using computerized algorithms.

#### Matrix-Assisted Laser Desorption Ionization Mass Spectrometry Imaging (MALDI MSI) for spatial distribution of N-glycans

Fresh-frozen brains were cut into sagittal sections with a thickness of 12 μm on a Leica CM3050 cryostat and then thaw-mounted on a conductive ITO slide (ASTA, Gyeonggi-do, Republic of Korea). The sample slides were dried under vacuum for 2 hours. The sample preparation for MALDI MSI of N-glycans was performed as described^45^, with minor modifications. Briefly, the dried sample slides were incubated at 60 °C for 1 hour and washed with xylene, ethanol, Carnoy’s solution (ethanol:chloroform:acetic acid=6:3:1), and DW. Next, 0.1 mg/ml PNGase F PRIME-LY™ ULTRA GLYCOSIDASE (N-zyme scientifics, PA, USA) was applied to the sample slide using an HTX M5 sprayer (HTX technologies, NC, USA). The sample slides were placed in a chamber maintained at 97% relative humidity using 150 g/L K2SO4 and incubated at 37 °C for 2 hours. Then, 7 mg/ml α-cyano-4-hydroxycinnamic acid (CHCA; Sigma, MO, USA) added to 50% acetonitrile was applied with an HTX M5 sprayer.

N-glycan images of the mouse brain were obtained using MALDI QTOF instrument, timsTOF fleX (Bruker Daltonik GmbH, Bremen, Germany). Mass spectra were recorded using 1,000 laser shots per each pixel in positive ion mode. The mass range was set to 500 – 3000 Da, and the laser focusing size and spatial resolution were set to 50 μm. Mass calibration was performed using ESI tuning mix (Agilent, CA, USA) and peptide calibration standard (Bruker Daltonik GmbH, Bremen, Germany). The measured mass spectra were converted into images in SCiLS Lab 2023a software (Bruker Daltonik GmbH, Bremen, Germany).

#### Protein enrichment from mouse brain tissue

The dried membrane fraction was mixed with 100 μL of 50 mM ammonium bicarbonate. After then, 2 μL of 550 mM dithiothreitol were added, and samples were incubated in a water bath at 60 °C for 50 min. The chilled samples mixed with 4 μL of 450 mM indole-3-acetic acid (IAA) were incubated at room temperature for 45 min in dark. The mixture was digested with 10 μL of 0.2 g/L trypsin in a water bath at 37 °C for 16 hours. The digested proteins were purified by C18 SPE. C18 cartridges (Thermo Fisher Scientific) were first washed with 6 mL of 0.1% TFA in DI water (v/v) and 6 mL of 80% ACN, 0.1% TFA in DI water (v/v). And the cartridge was conditioned with 6 mL of 0.1% TFA in DI water (v/v), followed by loading aqueous digested protein solution onto the cartridge. Continuously, peptides were eluted with 6 mL of 80% ACN, 0.1% TFA in DI water (v/v) after washing with 6 mL of 0.1% TFA in DI water (v/v). The fraction was dried under vacuum and resolubilized with 100 μL of 0.1% TFA in DI water (v/v) before LC-MS/MS analysis.

#### Proteome analysis with nano-LC/Orbitrap MS/MS

Proteins and glycoproteins were analyzed using a Thermo Scientific Ultimate 3000 RSLCnano system coupled to a Thermo Scientific Q Exactive Plus hybrid quadrupole Orbitrap mass spectrometer with a nano-electrospray ion source. Mobile phases A and B were water with 0.1% formic acid (v/v) and ACN with 0.1% formic acid (v/v), respectively. The samples were separated with 0.3 μL /min flow rate for 130 min using PepMap RSLC C18 column (Thermo Scientific, 2.0 μm, 75 μm × 50 cm). The Orbitrap MS parameters were set as follow: survey scan of peptide precursors was performed at 70 K FWHM resolution in the range of *m/z* 350-1900. HCD fragmentation was performed on 27 at a resolving power setting of 17.5 K. Resulting fragments detected in the range of *m/z* 200-2000 was used.

The raw data of proteins were processed using MaxQuant v2.0.1.0. Data were searched against the UniProt/SwissProt mouse (Mus musculus) protein database with 17,127 total entries and contaminant proteins. Searches were performed with the following parameter: Data were filtered with a peptide -to-spectrum match (PSM) of 0.01 false discovery rates (FDR), 7 in minimum peptide length, 1 in minimum unique peptide, modification including oxidation and acetylation in protein N-terminal, and true of iBAQ and match between runs. LFQ intensity data was statistically calculated concentrations of proteins using Perseus. Gene Ontology (GO) term enrichment was performed using PANTHER,db, and network clusters were by Cytoscape using ClueGO and CluePedia.

#### Statistical analysis of data

All data are presented as the mean ± SEM and significant symbol and value were represented in each figure and legend, respectively. GraphPad Prism 9.4.1 software was used for statistical analysis. Normal distribution was firstly assessed using the D’Agostino-Pearson omnibus normality test for all experiments. Parametric tests (Student’s two-tailed unpaired t-test, one-way ANOVA with Dunnett’s post hoc test) were used for data following normal distribution. Non-parametric tests (Mann Whitney test, Kruskal-Wallis test with Dunn’s post hoc test) were used for data not following normal distribution. Samples that passed normality test, but not equal variance test were assessed with Welch’s correction. Two-way ANOVA followed by Holm-Šídák’s post hoc test was used. The significance symbol was represented as asterisks (*p<0.05, **p<0.01, ***p<0.001, ****p<0.0001, and ns=non-significant).

## Acknowledgments

This study was supported by Center for Cognition and Sociality (IBS-R001-D2) to C.J.L., and a Young Scientist Fellowship (IBS-R001-Y1) to W.K. from the Institute for Basic Science (IBS), South Korea.

## References

1 Rivinoja, A., Hassinen, A., Kokkonen, N., Kauppila, A. & Kellokumpu, S. J. J. o. c. p. Elevated Golgi pH impairs terminal N-glycosylation by inducing mislocalization of Golgi glycosyltransferases. 220, 144–154 (2009).

2 Maeda, Y. & Kinoshita, T. J. M. i. e. The acidic environment of the Golgi is critical for glycosylation and transport. 480, 495–510 (2010).

3 Fan, J. et al. Golgi apparatus and neurodegenerative diseases. 26, 523–534 (2008).

4 Joshi, G., Bekier, M. E. & Wang, Y. J. F. i. n. Golgi fragmentation in Alzheimer’s disease. 9, 340 (2015).

5 Sundaramoorthy, V., Sultana, J. M. & Atkin, J. D. J. F. i. N. Golgi fragmentation in amyotrophic lateral sclerosis, an overview of possible triggers and consequences. 9, 400 (2015).

6 Kellokumpu, S. J. F. i. c. & biology, d. Golgi pH, ion and redox homeostasis: how much do they really matter? 7, 93 (2019).

7 Marshansky, V. & Futai, M. The V-type H+-ATPase in vesicular trafficking: targeting, regulation and function. Curr Opin Cell Biol 20, 415–426, doi:10.1016/j.ceb.2008.03.015 (2008).

8 Nakamura, N., Tanaka, S., Teko, Y., Mitsui, K. & Kanazawa, H. Four Na+/H+ exchanger isoforms are distributed to Golgi and post-Golgi compartments and are involved in organelle pH regulation. J Biol Chem 280, 1561–1572, doi:10.1074/jbc.M410041200 (2005).

9 Numata, M. & Orlowski, J. Molecular cloning and characterization of a novel (Na+,K+)/H+ exchanger localized to the trans-Golgi network. J Biol Chem 276, 17387–17394, doi:10.1074/jbc.M101319200 (2001).

10 Maeda, Y., Ide, T., Koike, M., Uchiyama, Y. & Kinoshita, T. GPHR is a novel anion channel critical for acidification and functions of the Golgi apparatus. Nat Cell Biol 10, 1135–1145, doi:10.1038/ncb1773 (2008).

11 Sou, Y. S. et al. Cerebellar Neurodegeneration and Neuronal Circuit Remodeling in Golgi pH Regulator-Deficient Mice. eNeuro 6, doi:10.1523/ENEURO.0427-18.2019 (2019).

12 Brunner, J. D., Lim, N. K., Schenck, S., Duerst, A. & Dutzler, R. X-ray structure of a calcium-activated TMEM16 lipid scramblase. Nature 516, 207–212, doi:10.1038/nature13984 (2014).

13 Caputo, A. et al. TMEM16A, a membrane protein associated with calcium-dependent chloride channel activity. Science 322, 590–594, doi:10.1126/science.1163518 (2008).

14 Dang, S. et al. Cryo-EM structures of the TMEM16A calcium-activated chloride channel. Nature 552, 426–429, doi:10.1038/nature25024 (2017).

15 Schroeder, B. C., Cheng, T., Jan, Y. N. & Jan, L. Y. Expression cloning of TMEM16A as a calcium-activated chloride channel subunit.>Cell 134, 1019–1029, doi:10.1016/j.cell.2008.09.003 (2008).

16 Yang, Y. D. et al. TMEM16A confers receptor-activated calcium-dependent chloride conductance. Nature 455, 1210–1215, doi:10.1038/nature07313 (2008).

17 Beaulieu-Laroche, L. et al. TACAN Is an Ion Channel Involved in Sensing Mechanical Pain. Cell 180, 956–967 e917, doi:10.1016/j.cell.2020.01.033 (2020).

18 Rong, Y. et al. TMEM120A contains a specific coenzyme A-binding site and might not mediate poking- or stretch-induced channel activities in cells. Elife 10, doi:10.7554/eLife.71474 (2021).

19 Lee, C. et al. The lysosomal potassium channel TMEM175 adopts a novel tetrameric architecture. Nature 547, 472–475, doi:10.1038/nature23269 (2017).

20 Cang, C., Aranda, K., Seo, Y. J., Gasnier, B. & Ren, D. TMEM175 Is an Organelle K(+) Channel Regulating Lysosomal Function. Cell 162, 1101–1112, doi:10.1016/j.cell.2015.08.002 (2015).

21 Yang, J. et al. PAC, an evolutionarily conserved membrane protein, is a proton-activated chloride channel. Science 364, 395–399, doi:10.1126/science.aav9739 (2019).

22 Hirata, T. et al. Post-Golgi anterograde transport requires GARP-dependent endosome-to-TGN retrograde transport. Mol Biol Cell 26, 3071–3084, doi:10.1091/mbc.E14-11-1568 (2015).

23 Shin, J. J. H. et al. Spatial proteomics defines the content of trafficking vesicles captured by golgin tethers. Nat Commun 11, 5987, doi:10.1038/s41467-020-19840-4 (2020).

24 Patkunarajah, A. et al. TMEM87a/Elkin1, a component of a novel mechanoelectrical transduction pathway, modulates melanoma adhesion and migration. Elife 9, doi:10.7554/eLife.53308 (2020).

25 Hoel, C. M., Zhang, L. & Brohawn, S. G. J. b. Structure of the GOLD-domain seven-transmembrane helix protein family member TMEM87A. (2022).

26 Doyle, D. A. et al. The structure of the potassium channel: molecular basis of K+ conduction and selectivity. Science 280, 69–77, doi:10.1126/science.280.5360.69 (1998).

27 Zhang, Y. et al. An RNA-sequencing transcriptome and splicing database of glia, neurons, and vascular cells of the cerebral cortex. J Neurosci 34, 11929–11947, doi:10.1523/JNEUROSCI.1860-14.2014 (2014).

28 Zhang, Y. et al. Purification and Characterization of Progenitor and Mature Human Astrocytes Reveals Transcriptional and Functional Differences with Mouse. Neuron 89, 37–53, doi:10.1016/j.neuron.2015.11.013 (2016).

29 Linders, P. T. A., Ioannidis, M., Ter Beest, M. & van den Bogaart, G. Fluorescence Lifetime Imaging of pH along the Secretory Pathway. ACS Chem Biol 17, 240–251, doi:10.1021/acschembio.1c00907 (2022).

30 Antoniel, M. et al. The unique histidine in OSCP subunit of F-ATP synthase mediates inhibition of the permeability transition pore by acidic pH. EMBO Rep 19, 257–268, doi:10.15252/embr.201744705 (2018).

31 Jumper, J. et al. Highly accurate protein structure prediction with AlphaFold. Nature 596, 583–589, doi:10.1038/s41586-021-03819-2 (2021).

32 Varadi, M. et al. AlphaFold Protein Structure Database: massively expanding the structural coverage of protein-sequence space with high-accuracy models. Nucleic Acids Res 50, D439–D444, doi:10.1093/nar/gkab1061 (2022).

33 Schapiro, F. B. & Grinstein, S. Determinants of the pH of the Golgi complex. J Biol Chem 275, 21025–21032, doi:10.1074/jbc.M002386200 (2000).

34 Matamala, E., Castillo, C., Vivar, J. P., Rojas, P. A. & Brauchi, S. E. Imaging the electrical activity of organelles in living cells. Commun Biol 4, 389, doi:10.1038/s42003-021-01916-6 (2021).

35 Lee, J. et al. Spatial and temporal diversity of glycome expression in mammalian brain. Proc Natl Acad Sci U S A 117, 28743–28753, doi:10.1073/pnas.2014207117 (2020).

36 Ji, I. J. et al. Spatially-resolved exploration of the mouse brain glycome by tissue glycocapture (TGC) and nano-LC/MS. Anal Chem 87, 2869–2877, doi:10.1021/ac504339t (2015).

37 Hodgkin, A. L. & Huxley, A. F. J. T. J. o. p. The dual effect of membrane potential on sodium conductance in the giant axon of Loligo. 116, 497 (1952).

38 Buettner, R., Papoutsoglou, G., Scemes, E., Spray, D. C. & Dermietzel, R. Evidence for secretory pathway localization of a voltage-dependent anion channel isoform. Proc Natl Acad Sci U S A 97, 3201–3206, doi:10.1073/pnas.97.7.3201 (2000).

39 Gupta, R. & Ghosh, S. Phosphorylation of purified mitochondrial Voltage-Dependent Anion Channel by c-Jun N-terminal Kinase-3 modifies channel voltage-dependence. Bíochím Open 4, 78–87, doi:10.1016/j.biopen.2017.03.002 (2017).

40 Ju, Y. H. et al. Astrocytic urea cycle detoxifies Abeta-derived ammonia while impairing memory in Alzheimer’s disease. Cell Metab 34, 1104–1120 e1108, doi:10.1016/j.cmet.2022.05.011 (2022).

41 Ma, L., Ouyang, Q., Werthmann, G. C., Thompson, H. M. & Morrow, E. M. Live-cell Microscopy and Fluorescence-based Measurement of Luminal pH in Intracellular Organelles. Front Cell Dev Biol 5, 71, doi:10.3389/fcell.2017.00071 (2017).

42 Kwak, H. et al. Astrocytes Control Sensory Acuity via Tonic Inhibition in the Thalamus. Neuron 108, 691–706 e610, doi:10.1016/j.neuron.2020.08.013 (2020).

43 Cruz-Sanchez, A. et al. Developmental onset distinguishes three types of spontaneous recognition memory in mice. Sci Rep 10, 10612, doi:10.1038/s41598-020-67619-w (2020).

44 An, H. J. et al. Extensive determination of glycan heterogeneity reveals an unusual abundance of high mannose glycans in enriched plasma membranes of human embryonic stem cells. Mol Cell Proteomics 11, M111 010660, doi:10.1074/mcp.M111.010660 (2012).

45 Drake, R. R., Powers, T. W., Norris-Caneda, K., Mehta, A. S. & Angel, P. M. In Situ Imaging of N-Glycans by MALDI Imaging Mass Spectrometry of Fresh or Formalin-Fixed Paraffin-Embedded Tissue. Curr Protoc Protein Sci 94, e68, doi:10.1002/cpps.68 (2018).

