## Extended Data Figure for "GolpHCat (TMEM87A): a unique voltage-gated and pH-sensitive cation channel in the Golgi"

**Extended Data Figure 1**

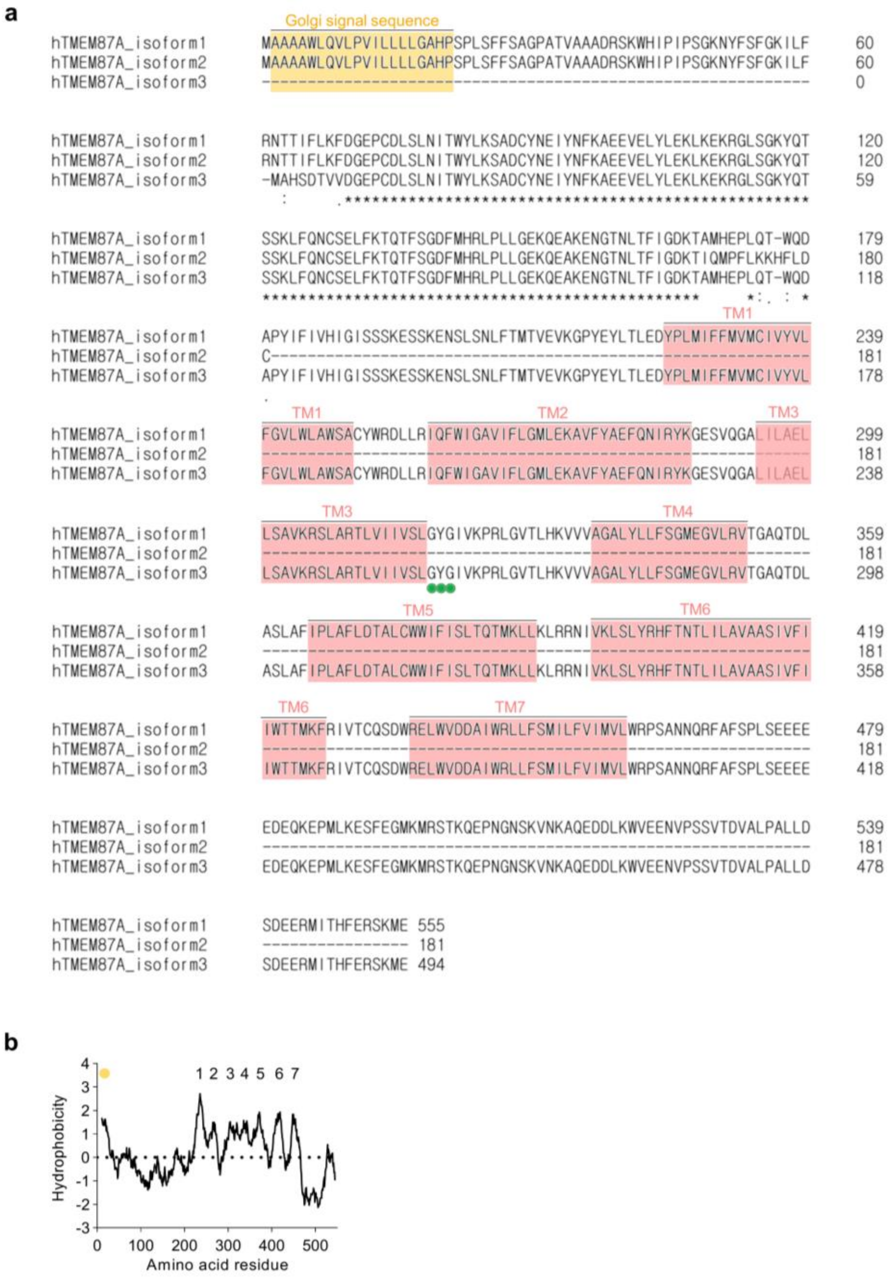


**Extended Data Fig. 1 Amino acid sequence alignment of human TMEM87A isoforms.**
**a,** Multiple protein sequence alignment for human TMEM87A isoform 1, 2 and 3 generated with the Clustal-Omega. Yellow and pink boxes indicate signal sequence and TMs, respectively. Green circles indicate GYG motif sequence. **b,** Hydrophobicity plot of hTMEM87A. Yellow circle indicates a predicted Golgi signal sequence site.

**Extended Data Figure 2**

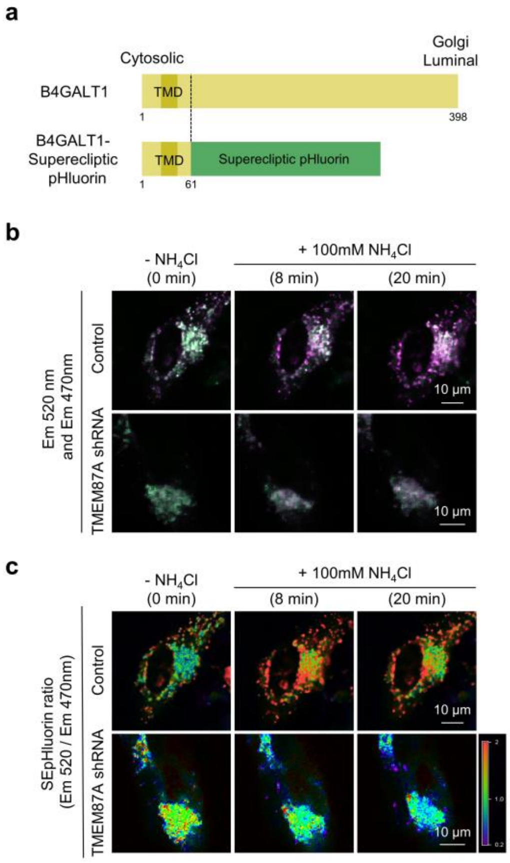


**Extended Data Fig. 2 TMEM87A is necessary for Golgi pH homeostasis.**
**a,** Full length of B4GALT1 (top). Constructs of B4GALT1 TMD-fused SEpHluorin to target SEpHluorin in the Golgi lumen (bottom). **b,** Representative merged fluorescence images illuminated with two emissions 470nm and 520nm at 100 mM NH4Cl treat before and after in Control and TMEM87A shRNA transfected cultured human astrocytes. **c,** Representative fluorescence ratio (em 520nm/em 470nm) images.

**Extended Data Figure 3**

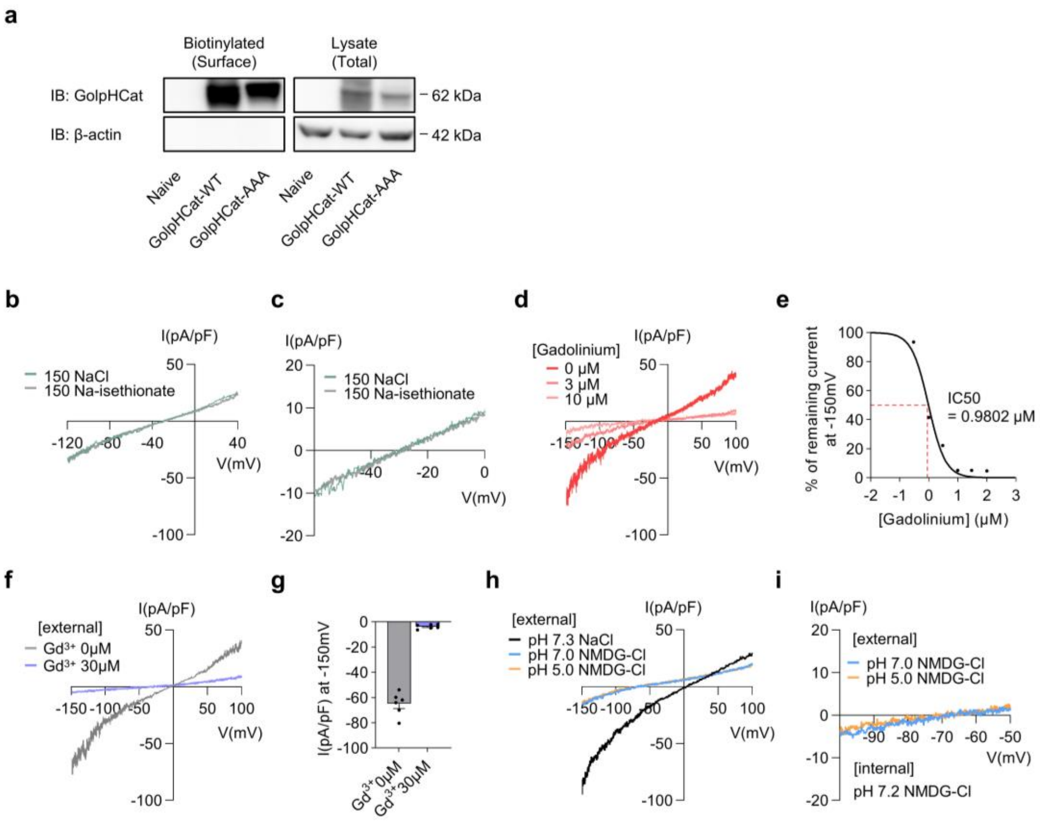


**Extended Data Fig. 3 TMEM87A mediates currents carried by cation and blocked by non-selective channel blocker in heterologous system.**

**a,** Surface biotinylation assay of TMEM87A WT and TMEM87A-AAA transfected cells blotted with TMEM87A antibody. **b,** Representative I-V relationship from TMEM87A WT transfected cells under replacing the bath solution NaCl to Na-isethionate. The currents were corrected by liquid junction potential (LJP), respectively. (NaCl, green; Na-isethionate, gray; n=3). **c,** Magnified trace for reversal potential from (**b**)**. d,** Representative I-V relationship from TMEM87A WT transfected cells under the bath solution containing gadolinium. **e,** Dose-response curve for percentage currents at -150 mV for gadolinium (0, 0.3, 1, 3, 10, 30 and 100 µM) (n=5). **f,** Representative I-V relationship from TMEM87A WT transfected cells under the bath solution without ($Gd^{3+}$ 0 µM) or with 30 µM gadolinium ($Gd^{3+}$ 30 µM). **g,** Current densities measured at -150mV from (**f**); ($Gd^{3+}$ 0 µM vs. $Gd^{3+}$ 30 µM, $p<0.0001$; n=6). **h,** Representative I-V relationship from TMEM87A WT transfected cells under replacing the bath solution pH 7.3 NaCl to pH 7.0 and 5.0 NMDG-Cl. (pH 7.3 NaCl, black; pH 7.0 NMDG-Cl, blue; pH 5.0 NMDG-Cl, orange; n=3). **i,** Magnified trace for reversal potential from (**h**)**.**

**Extended Data Figure 4**

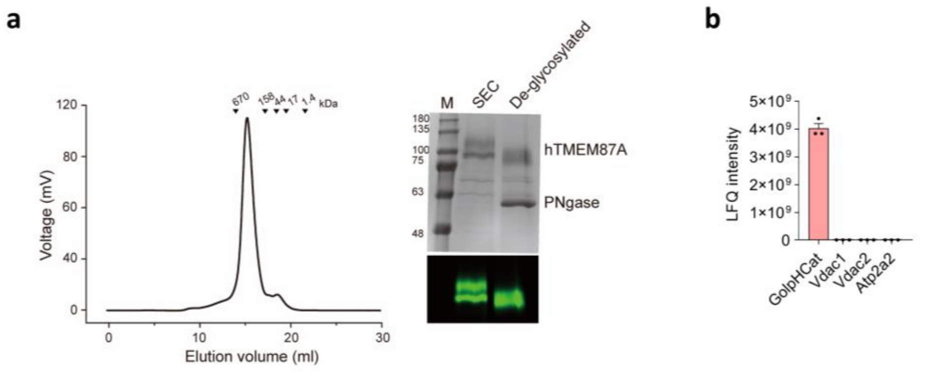


**Extended Data Fig. 4 FSEC profiles of EGFP-tagged TMEM87A.**

**a,** Size-exclusion chromatography (SEC) profile of hTMEM87A-EGFP-Twin-strep (left). SDS-PAGE analysis of hTMEM87A-EGFP-Twin-strep after PNGase F treatment (right). **b,** bar graph of LFQ intensity of GolpHCat and other pore-forming ion channels involved in TMEM87A purified solution.

**Extended Data Figure 5**

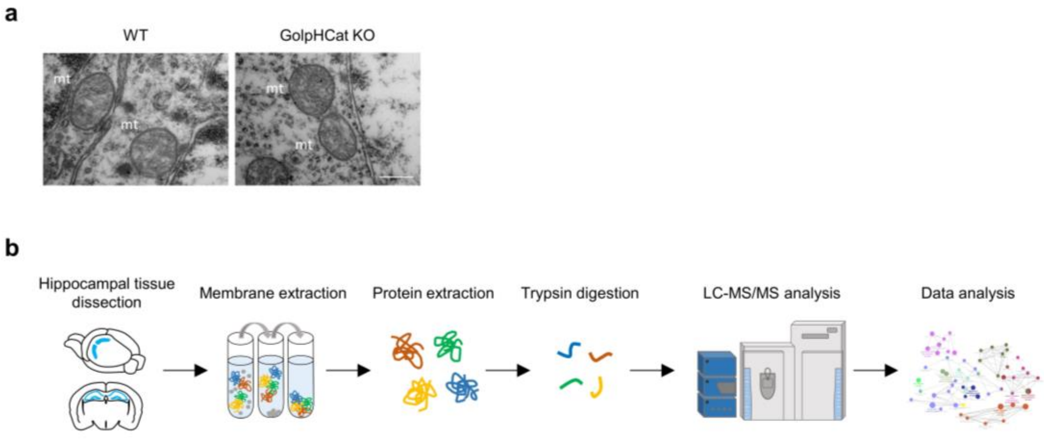


**Extended Data Fig. 5**

**a,** Representative TEM images of the mitochondria in WT and GolpHCat KO mice. Scale bar, 500nm. **b,** Workflow of proteomics analysis with hippocampus sample.

**Extended Data Figure 6**

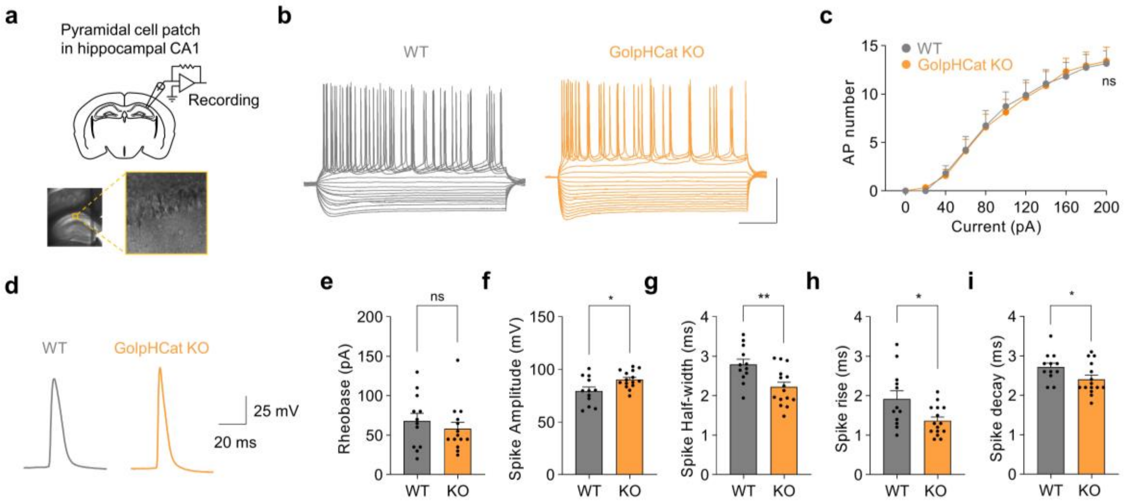


**Extended Data Fig. 6 Intrinsic neuronal excitability of the CA1 pyramidal neurons in WT and GolpHCat KO mice.**

**a,** Schematic diagram of a slice patch from WT or GolpHCat KO mice (top) and magnified differential interference contrast (DIC) image of a whole-cell patch-clamped CA1 pyramidal neuron in hippocampus (bottom). **b,** Representative action potential traces in the CA1 pyramidal neurons in WT and GolpHCat KO mice. Scale bar: 50 mV and 200 ms. **c,** Spike (action potential) number under the depolarizing current step protocol from 0 pA to +200 pA (in 20 pA increments) in the CA1 pyramidal neurons in WT (n=12, 3 mice) and GolpHCat KO (n=15, 3 mice) mice; (WT vs. GolpHCat KO, p=0.9838). **d,** Representative action potential traces of WT and GolpHCat KO mice. **e,** Rheobase; (WT vs. GolpHCat KO, p=0.4683). **f,** Rheobase spike amplitude; (WT vs. GolpHCat KO, p=0.0143). **g,** Rheobase spike half-width; (WT vs. GolpHCat KO, p=0.0042). **h,** Rheobase spike rise; (WT vs. GolpHCat KO, p=0.0304). **i,** Rheobase spike decay; (WT vs. GolpHCat KO, p=0.0464). Data are presented as the mean ± SEM. *p<0.05, **p<0.01, ns=non-significant. Two-way ANOVA followed by Holm-Šídák's multiple comparisons test in **(c)**, Mann-Whitney test in (**e**), Unpaired t-test in (**f,g,i**), Unpaired t-test with Welch's correction (**h**).

**Extended Data Figure 7**

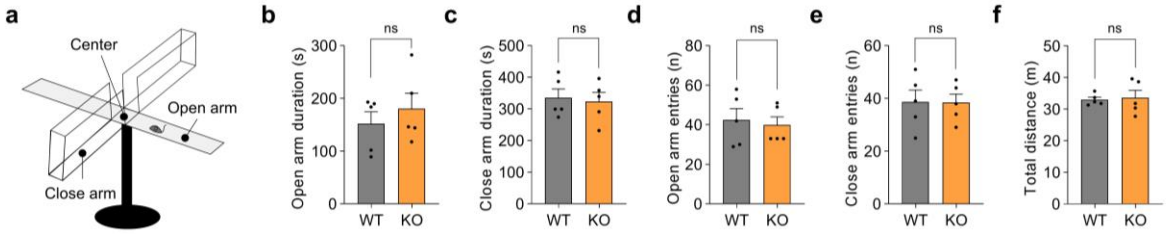


**Extended Data Fig. 7 Anxiety level was not altered in GolpHCat KO mice.**
**a,** Schematic diagram of the elevated plus maze (EPM). **b-f,** bar graph of open arm duration (**b**), close arm duration (**c**), open arm entries (**d**), close arm entries (**e**), and total distance (**f**) in elevated plus maze test of WT (n=5) and GolpHCat KO (n=5) mice. Data are presented as the mean ± SEM. ns=non-significant. Mann-Whitney test in (**b-f**).

**Extended Data Figure 8**

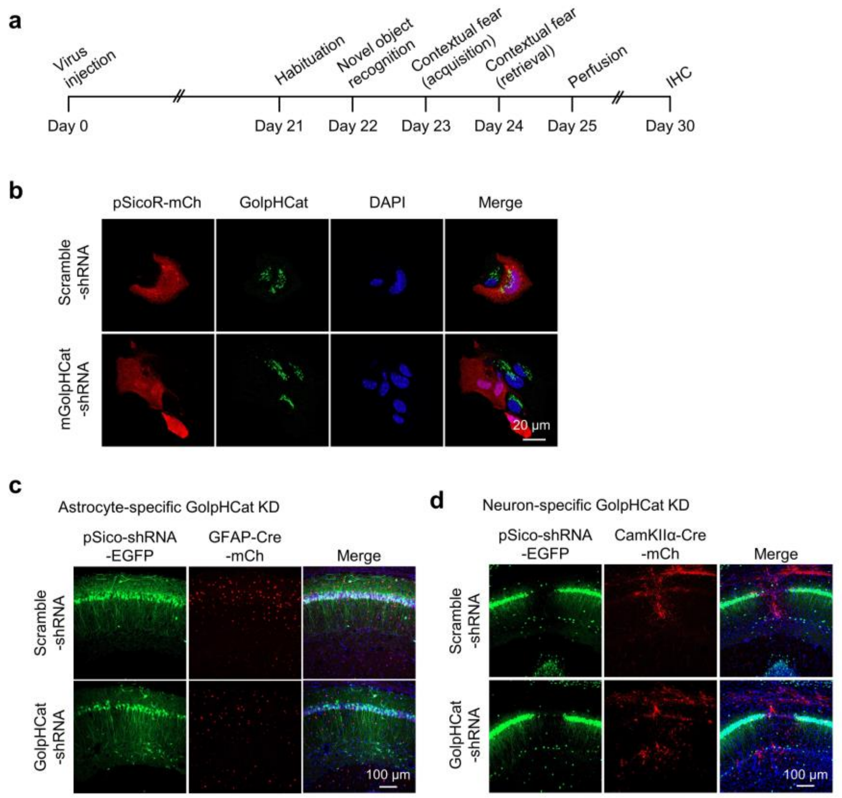


**Extended Data Fig. 8 Genetic ablation of GolpHCat of astrocytes and neurons in hippocampus.**
**a,** Experimental timeline for virus injection, behavior tests, and IHC. **b.** Immunostaining of GolpHCat in the scramble shRNA or TMEM87A shRNA microporated cultured hippocampal astrocytes for confirming the efficiency of GolpHCat shRNA. Scare bar, 20 µm. **c**, Fluorescence images in AAV-GFAP-cre-mCh + Lenti-pSico-Scrambled / GolpHCat shRNA-GFP virus-injected mice. Scare bar, 100 µm. **d**, Fluorescence images in AAV-CamKII-cre-mCh + Lenti-pSico-Scrambled / GolpHCat shRNA-GFP virus-injected mice. Scare bar, 100 µm.
